# Pili-like proteins are essential for conjugation also in Gram-positive bacteria

**DOI:** 10.64898/2025.12.18.695078

**Authors:** Josy ter Beek, Dennis Svedberg, Kieran Deane-Alder, May I.A. Ibrahim, André Mateus, Ronnie P.-A. Berntsson

## Abstract

Type IV secretion systems (T4SS) enable the spread of antibiotic resistance and other virulence factors. In Gram-positive bacteria, T4SSs have long been thought to lack VirB2-like proteins that form conjugative pili and instead rely on adhesins for cell-cell contacts. Yet, it has remained unclear how subsequent DNA transfer (conjugation) is mediated. Here we identify a VirB2-like protein, PrgF_B2_, from the clinically isolated plasmid pCF10 in *Enterococcus faecalis* and show that it is essential for conjugation. Structural modelling confidently predicts a pilus-like assembly for PrgF_B2_. This is in line with recently proposed models for conjugative pili in Gram-negative T4SSs. We validate the PrgF_B2_ pili prediction through mutagenesis, conjugation assays, and targeted chemical labelling. By combining various machine learning bioinformatic techniques, we analysed >1000 Gram-positive conjugative plasmids from diverse species, including major pathogens, and identified probable pili-forming VirB2-like proteins in a large majority. Our findings challenge the prevailing view that Gram-positive T4SSs lack pili and highlight critical targets for combating antimicrobial resistance and virulence in Gram-positive bacteria.

## Introduction

Type 4 secretion systems (T4SSs) are megadalton-sized membrane protein complexes that facilitate the transfer of proteins and/or DNA from bacterial donor cells to recipient cells. Conjugative T4SSs are main drivers for horizontal DNA transfer in bacteria and are of major interest as they spread antibiotic resistance and other virulence factors. T4SSs have been studied for decades, and multiple labs have provided structural and functional models for these systems, but the focus has been on a few model systems from Gram-negative (G-) bacteria(Paillard *et al*, 2025a).

In conjugative T4SSs, the mobile DNA, often encoded on a plasmid, is recognized by its *oriT* sequence. A specific relaxase then nicks the DNA, makes it single stranded and becomes covalently attached to the 5’ end. With the help of the coupling protein (T4CP), this protein:ssDNA substrate is then translocated through the T4SS channel. In G-T4SSs the transport channel comprises an inner membrane complex, an outer membrane complex, a connecting stalk and a conjugative pilus that extends from the cell. Pili of G-T4SSs mainly consist of helical polymers of VirB2-like proteins. Although it has been debated whether these pili form the transport channel for the protein:ssDNA substrate, this has been shown for the conjugative T4SSs from the F and pED208 plasmids from *E. coli (Goldlust et al, 2023; Beltrán et al, 2024; Babić et al, 2008)*.

Pili forming VirB2-like proteins are essential in G-T4SSs (Goldlust *et al*, 2023; Vadakkepat *et al*, 2024a). Despite high sequence divergence, all characterized mature VirB2 homologs are small membrane proteins with two transmembrane helices, connected by a short and charged cytoplasmic loop (Vadakkepat *et al*, 2024b; Paiva *et al*, 1992; Beijersbergen *et al*, 1994). Pili assembly is presumed to occur from a pool of these membrane-inserted VirB2 proteins that oligomerize onto the VirB6 pentamer and thereby get extruded from the membrane (Macé *et al*, 2022; Paillard *et al*, 2025a). The VirB6 pentamer was until recently seen as the central inner membrane complex of the T4SS, but two separate groups have now presented findings that revise this model(Paillard *et al*, 2026; Bloch *et al*, 2026). VirB2 is instead proposed to during the assembly process push out the VirB6 pentamer from the central T4SS complex (Fig. 1A)^10,11^. In the newly presented cryo-electron microscopy structures, VirB6 is located in between the long VirB2 oligomeric polymer and the VirB5 pentamer at the tip that facilitates adhesion to recipient cells. In line with other available structures of assembled conjugative pili, the charged loop of the VirB2 proteins faces the lumen, while the N and C-terminal ends are on the outside of the tubular structure. The charge of the lumen is often modulated by specifically bound phospholipids (Tiago Costa *et al*, 2016; Amro *et al*, 2023; Vadakkepat *et al*, 2024b; Ishimoto *et al*, 2025b).

**Figure 1.**
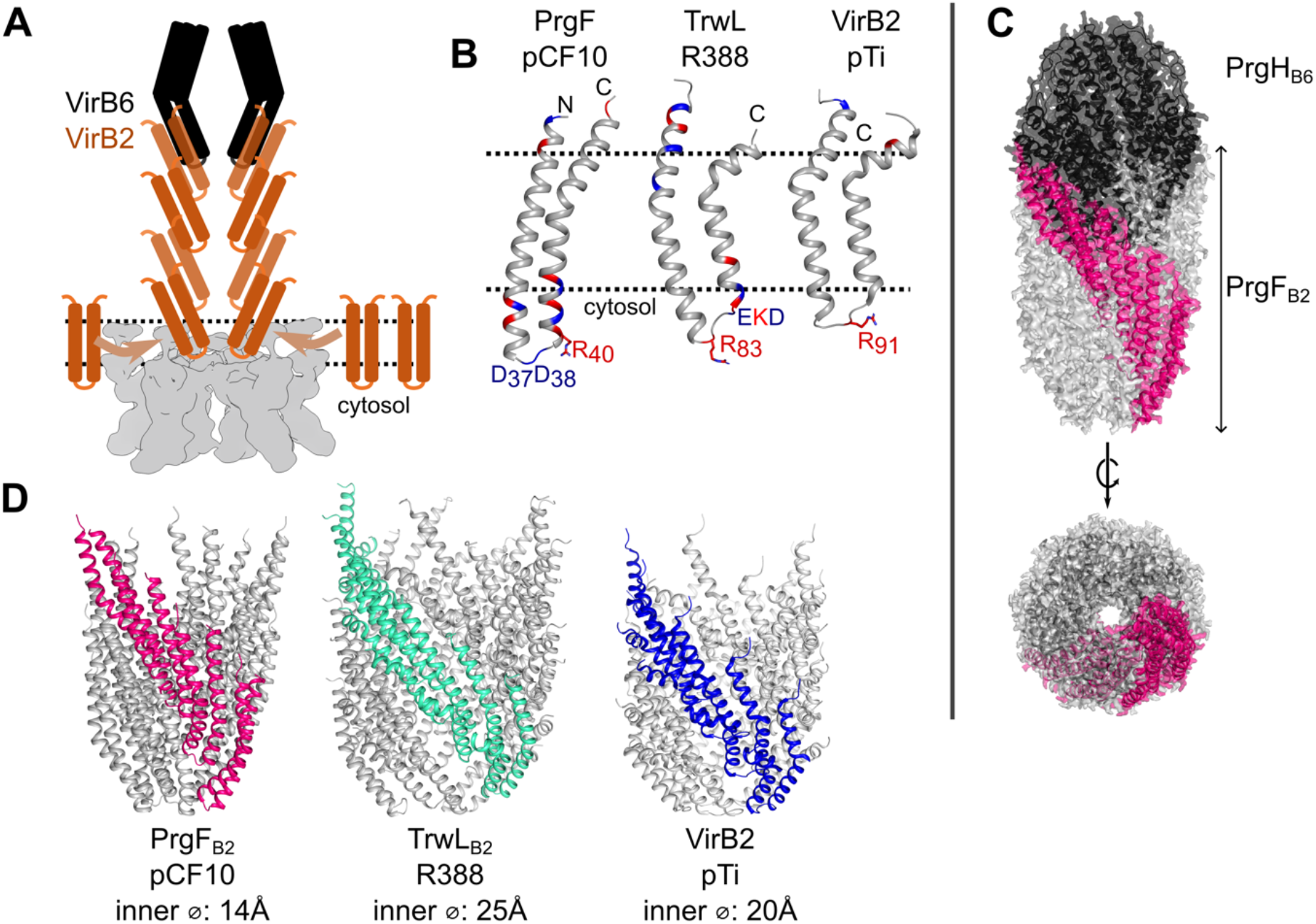
Overview of VirB2-like pili forming proteins. A) Cartoon visualizing the membrane insertion and assembly model for VirB2-like proteins (brown) into a pilus in the T4SS channel, thereby pushing the VirB6 (black) pentamer outwards (slice-through showing only 2 copies of the pentamer). The hexameric VirB4 ATPase is rendered in grey underneath. Figure adapted from Paillard *et al*, 2026 and Bloch *et al*, 2026 (Paillard *et al*, 2026; Bloch *et al*, 2026). B) AlphaFold model of PrgF alongside characterized VirB2-like proteins from R388 (PDB ID: 8S6H) and pTi (PDB ID: 8CUE). Dotted lines indicate the approximate location of the membrane. C) Overview of a PrgF_B2_-PrgH_B6_ AlphaFold model, with the PrgH_B6_ pentamer shown in black, 4 copies of PrgF_B2_ visualizing the helical rise in pink and the remaining 16 copies of PrgF_B2_ in grey. D) Comparison of the PrgF_B2_ pili model (as in C, but without showing PrgH_B6_) with the determined structures of the pili from R388 and pTi (PDB codes as in B).

We know much less of Gram-positive (G+) T4SSs, but they are consistently described to lack conjugative pili (Guglielmini *et al*, 2014; Minny Bhatty *et al*, 2013; Elisabeth Grohmann *et al*, 2003; Maffo-Woulefack *et al*, 2025; Ryan *et al*, 2023; Grohmann *et al*, 2018). It is known that some G+ T4SSs instead rely on cell-wall anchored adhesins for cell-to-cell contacts that enhance mating pair formation (Bhatty *et al*, 2015; Järvå *et al*, 2020; Revitt-Mills *et al*, 2021). These adhesins are important for conjugation to occur, but they are not essential and they do not form a channel for substrate transfer (Bhatty *et al*, 2015; Sun *et al*, 2023a; Revitt-Mills *et al*, 2021). Instead, different studies have suggested that such a channel could be formed either by the VirB1-like cell-wall hydrolases or the VirB8-like proteins in G+ T4SSs (Minny Bhatty *et al*, 2013; Maffo-Woulefack *et al*, 2025; Goessweiner-Mohr *et al*, 2014). However, all these proteins are membrane anchored and it is unclear how they could extend far enough to form a transfer pathway (Sun *et al*, 2024). How G+ T4SSs facilitate the transfer of the protein:ssDNA substrate through the cell-walls of both donor and recipient cells, which in *e.g.* enterococci equals roughly 80 nm, and then through the membrane of the recipient cell has therefore remained an unanswered question.

Despite all reports that G+ T4SS lack pili, we wondered whether they could have been overlooked, since identification of VirB2-like proteins is difficult as *virB2* genes are small and poorly conserved (Paillard *et al*, 2025a).

Here, we identify PrgF from the pheromone-inducible conjugative plasmid pCF10 of *Enterococcus faecalis* as a probable VirB2-like protein and show that it is essential for conjugation. AlphaFold modelling, mutagenesis and targeted chemical labelling support a model in which PrgF forms a conjugative pilus together with the VirB6-like protein PrgH_B6_. By combining various machine-learning bioinformatic techniques, we identify potential VirB2-like proteins in the majority of Gram-positive conjugative T4SSs, including systems from clinically relevant species.

## Results

### Model of a Gram-positive conjugative pilus

We previously modelled each predicted protein of the T4SS encoding *prgQ* operon from the pCF10 plasmid in *Enterococcus faecalis* (Breidenstein *et al*, 2025). One of these modelled proteins was PrgF, a small uncharacterized protein that is predicted to have two transmembrane helices (TMHs) connected by a short, charged loop that is predicted to be in the cytoplasm (Fig. S1). Its predicted structure bears some resemblance to the determined structures of mature VirB2-like proteins from G-bacteria (Fig. 1B), which are described to have the same orientation in the membrane (Fig. S1) (Beijersbergen *et al*, 1994; Paillard *et al*, 2025b).

AlphaFold modelling of oligomeric structures of PrgF only yielded low-confidence structures (Table S1). However, when modelling 5 or more PrgF subunits together with the pentameric VirB6-like protein from pCF10, PrgH_B6_, AlphaFold yielded high-confidence predictions of pilus-like structures (Fig. 1C-D, Fig. S2, Table S1). These models have ipTM scores of up to 0.87 (Table S1), show numerous interactions between PrgF and PrgH_B6_ (Fig. S3A) and have a helical architecture analogous to that observed in G-conjugative pili (Fig. 1D). To verify that PrgH_B6_ is not predicted to interact with other 2 TMH proteins, such as PrgI_B3_ or PcfH also from the pCF10 plasmid, we also modelled these potential complexes. This only gave low confidence models (ipTM < 0.45) that did not resemble a pilus (Table S2 and Fig. S3B), thus further strengthening the PrgF-PrgH_B6_ models.

It is noteworthy to mention that known VirB2-like pili-forming proteins from characterized G-T4SSs show the same behaviour as PrgF when being modelled with AlphaFold. Although all structures of conjugative pili currently available in the PDB contain just VirB2 oligomers, oligomeric AlphaFold models of these VirB2 proteins have low scores and do not resemble the determined structures, unless they are modelled together with a pentamer of their respective VirB6 (Table S3 and extended data table 1).

Based on these findings we hypothesized that PrgF might function as a VirB2-like protein and will therefore name it PrgF_B2_.

### The PrgFB2 membrane protein is essential for conjugation

To determine whether PrgF_B2_ is important for T4SS function, we deleted this gene from pCF10 in *E. faecalis* OG1RF. We kept 72 base pairs of *prgF*_B2_ in place to prevent changing the expression levels of downstream genes in the large pQ operon, which indeed remained unaltered (Fig. S4A-B). The deletion of *prgF*_B2_ resulted in a complete loss of conjugation efficiency (Fig. 2A and Fig. S5). Notably, conjugation was fully restored to wild-type levels upon complementation with heterologous expressed PrgF_B2_ from the pMSP:*prgF*_B2_ plasmid (Fig. 2A, blue bar). These findings demonstrate that, consistent with its proposed role as a VirB2-like protein, PrgF_B2_ is essential for conjugation.

**Figure 2.**
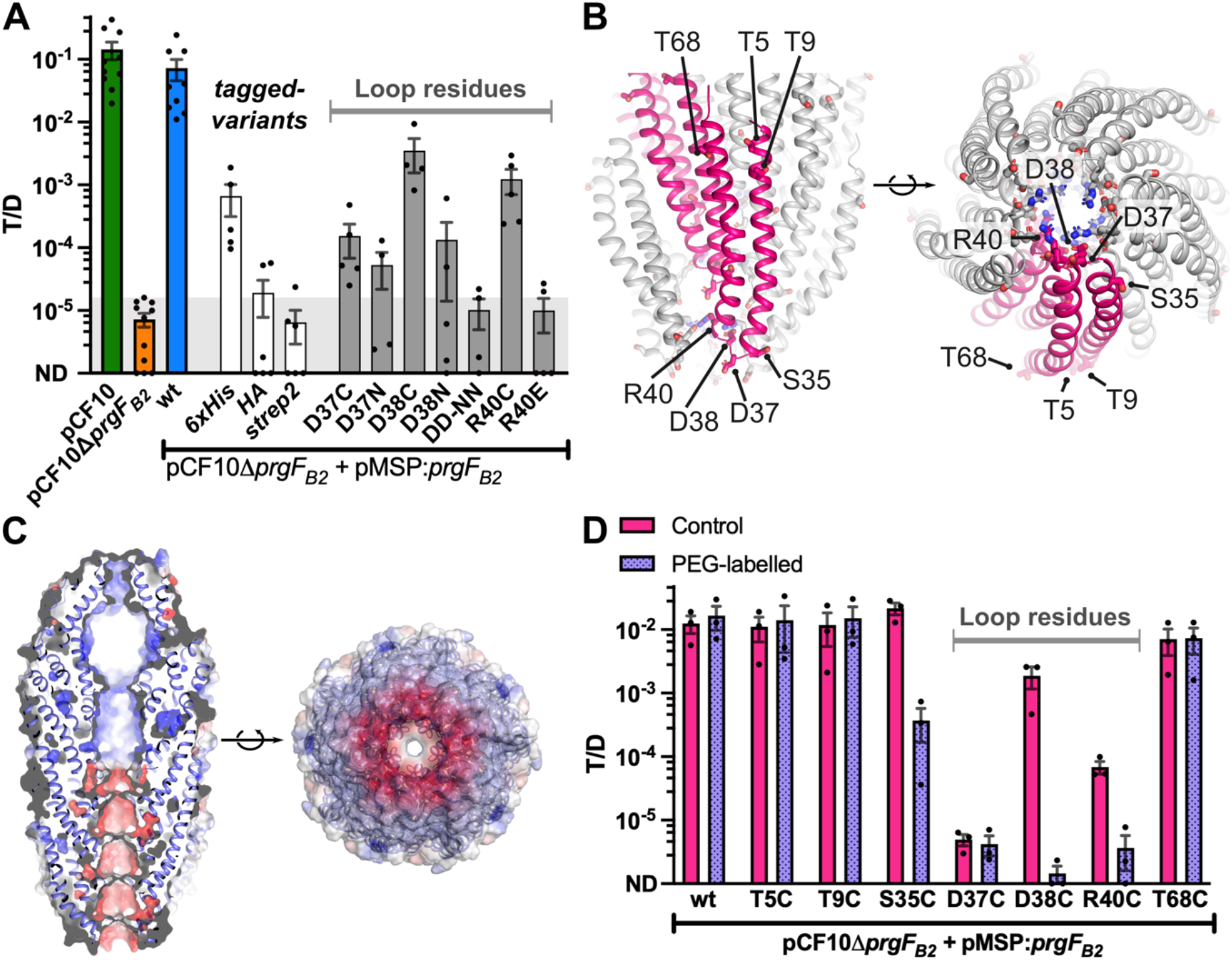
PrgF_B2_ is essential for conjugation and as in other VirB2-like proteins its charged loop is important for function. A) The conjugation efficiency of wild-type pCF10 (green bar) is completely reduced to background (indicated by shaded area, see Fig. S5) when *prgF*_B2_ is knocked-out (pCF10Δ*prgF*_B2_, orange bar). Conjugation is restored when wild-type PrgF_B2_ is expressed from a pMSP vector (blue bar). The other bars show pCF10Δ*prgF*_B2_ complemented with C-terminally tagged variants of PrgF_B2_ (white bars) or PrgF_B2_ variants with substituted loop residues (grey bars) all expressed from the pMSP vector. In the DD-NN variant both aspartates, D37 and D38, are exchanged for asparagine. Results without detected transconjugants were given a very small T/D score of 10^-6^ to take them along in the analysis and are shown as dots on the x-axis, ND = not detected. B) AlphaFold model of the PrgF_B2_ pili (as in Fig 1C-D) with the substituted residues shown as sticks and with their locations indicated for the upper pink subunit. C) Surface charge of the PrgF_B2_-PrgH_B6_ model (same as in Fig. 1C), showing a slice-through to visualize the lumen of the pili and a bottom (PrgF_B2_-side) view, coloured from red (-5 kT/e) to blue (+5 kT/e). D) Conjugation efficiency of PrgF_B2_ cysteine variants or wild type (wt) expressed from the pMSP vector in the pCF10Δ*prgF*_B2_ background without (control, in pink) or with maleimide-PEG_11_-biotin labelling (PEG-labelled, in purple and patterned). Results are shown as in panel A.

Tags were added to pMSP:*prgF*_B2_ to be able to probe expression and cellular localization via Western blot. Out of the tags tested, only a C-terminal 6xHis-tag retained some activity (Fig. 2A). Detection was challenging, probably due to very low expression levels of PrgF _B2_-His, its hydrophobic nature and small size (8.6 kDa). We were unable to detect PrgF_B2_-His in whole cells or soluble protein fractions but could detect a weak signal for PrgF_B2_-His in isolated membranes. Enrichment of PrgF_B2_-His by detergent solubilization, IMAC pull-down and further concentration greatly improved detection by Western blot (Fig. S4C). Concentrated fractions consistently showed an additional band ∼ 50 kDa even after denaturing SDS-PAGE, which might correspond to a pentamer of PrgF_B2_-His.

### Cytosolic loop residues are important for the function of PrgFB2

The AlphaFold model of the pilus suggests that D37, D38 and R40 from the charged loop of PrgF_B2_ line the pili lumen (Fig. 2B). In the structures from G-conjugative pili, the charged loop also lines the lumen with a bound phospholipid that modulates the overall surface charge of the pili cavity (Vadakkepat *et al*, 2024b). The surface charge of the PrgF_B2_-PrgH_B6_ AlphaFold model indicates that the initial cavity, formed by PrgH_B6_, has an overall positive surface charge, while the lumen of the PrgF_B2_ pilus is mainly negatively charged (Fig. 2C). In line with having an important functional role, we found that all modifications of charged residues in the PrgF_B2_ loop strongly reduced conjugation (Fig. 2A, grey bars). PrgF_B2_ variants where the charge of the R40 was reversed, or where both aspartates were exchanged for asparagine (Fig. 2A, DD-NN), could no longer support conjugation.

### PEG-labelling of residues predicted to point towards the pili lumen blocks conjugation

To further experimentally verify the AlphaFold model, we identified residues on the inside and outside of the proposed PrgF_B2_ pili-structure (Fig. 2B), exchanged them individually for cysteines, and used them for PEG-maleimide labelling after PrgF_B2_ expression in the presence of the rest of the T4SS (Ellison *et al*, 2019). We reasoned that the conjugation efficiency should be decreased if steric hindrances were introduced inside the lumen of the proposed PrgF_B2_ formed pili. To only probe transport with labelled PrgF_B2_, both pMSP:*prgF*_B2_ and the rest of the T4SS from the pQ operon (Fig. S4) on pCF10Δ*prgF*_B2_ were induced by addition of nisin and cCF10 (the inducing pheromone), respectively. Both inducers were washed away before labelling the donor cells for 10 minutes, directly followed by a very short (15 minutes) mating step with recipient cells. The recipient cell culture will contain naturally produced cCF10, but the removal of nisin stops *prgF*_B2_ transcription. With bulk mRNA half-life times being ca 5 minutes (Bryan *et al*, 2000; Laalami *et al*, 2014), only a very small amount of newly expressed PrgF_B2_ is expected to be present after the washing and labelling of the cells. The cell preparation procedure or the reduced mating time (from 2.5 hours to just 15 minutes), led to approximately 10-fold lower conjugation efficiencies (compare Fig. 2A & D).

Wild-type PrgF_B2_ does not contain any cysteines and is therefore not affected by maleimide-PEG_11_-biotin labelling. N-or C-terminal PrgF_B2_ residues that are predicted to be on the outside of the pili in the AlphaFold model, T5C, T9C, and T68C, have a conjugation efficiency like wild type and were also not affected by labelling. S35C, predicted to be in a protein-protein interface area of the pilus was as efficient in conjugation, but was partially affected by labelling. The residues situated in the charged loop that lines the pilus lumen in the model, D38C and R40C, showed a complete loss of conjugation efficiency after labelling (Fig. 2D). D37C, the last charged residue that we modified from the loop, has a very low conjugation efficiency making it difficult to determine if labelling affects the conjugation efficiency of this variant (Fig. 2D).

Addition of the maleimide-PEG11-biotin label did not affect conjugation with wild-type PrgF_B2_ or donor cell survival (Fig. S6), strongly indicating that the label does not enter cells as also described by the manufacturer. As in G-VirB2-like proteins, the loop of PrgF _B2_ is predicted to face the cytoplasm when PrgF_B2_ is in the membrane (Fig. 1B, S1) and loop-residues are therefore expected to only become accessible for labelling if PrgF_B2_ forms pili (Fig. 1A). Therefore, these results support a model in which PrgF_B2_ forms a pilus with loop residues D38 and R40 lining the lumen, consistent with the AlphaFold models.

### VirB2-like proteins are prevalent in G+ T4SSs, including many clinically relevant species

While assessing the function of PrgF_B2_, we wondered how widespread VirB2-like proteins are in G+ T4SSs. Therefore, we retrieved all annotated G+ conjugative plasmids from the plasmid database (PLSDB, >1000 plasmids) and clustered them based on their VirB6-like protein sequence similarity. We then selected 186 plasmids from a wide variety of G+ bacteria for further analysis (Fig. S7, Extended data 1).

To start our analysis, we took pCF10 and 13 other reference plasmids (representing plasmids that have previously been studied, from various clades of the representative clusters, Fig. S7). In these reference plasmids we searched all open reading frames to identify small proteins (50-170 amino acids) with 2 predicted TMH and with or without a signal peptide by DeepTMHMM analysis (Hallgren *et al*, 2022). This selection gives multiple candidates per plasmid and includes VirB3-like proteins. All selected proteins were further analysed by AlphaFold modelling, using 10 copies of the 2 TMH protein and 5 copies of the VirB6-like protein from the respective plasmid. The resulting AlphaFold models were subsequently analysed by their pTM, ipTM and PAE scores as well as by visual inspection. For all the 14 plasmids, only one of the candidates in each plasmid modelled as a pilus with an ipTM score separated from the other models (Fig. 3A). These will therefore further be referred to as the VirB2-like candidate models, while the other 2 TMH protein models will further be regarded as negative controls.

**Figure 3:**
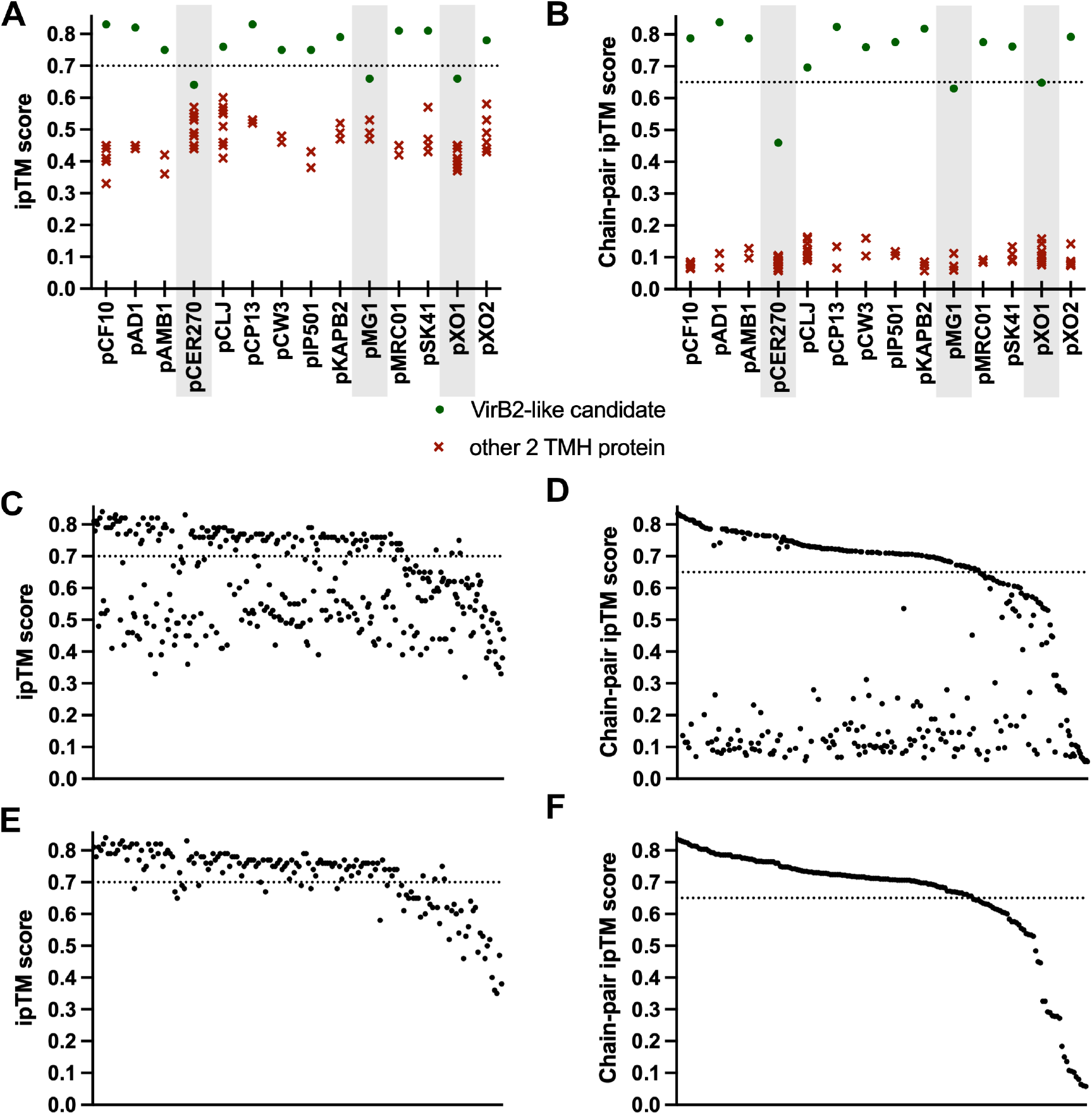
ipTM scores of AlphaFold models for 5 VirB6-like proteins (trimmed at the C-terminus as described in the materials and methods) and ten copies of the 2 TMH candidate protein. Chosen cut-off scores for reliable models are indicated by a dotted line. A-B) ipTM and chain-pair ipTM scores, respectively, for the 14 reference plasmids, with the identified complexes of VirB6- and VirB2-like proteins as green circles and the other investigated complexes as red crosses. C-D) ipTM and chain-pair ipTM scores for all AlphaFold models of all potential VirB2-like candidates with their VirB6-like protein from 172 representative plasmids. E-F) ipTM and chain-pair ipTM scores of only the best scoring model for each of the 172 plasmids.

As the VirB6 pentamer always predicts well, even our negative control models (Fig. 3A, S3, Table S2) have ipTM scores > 0.3. The accuracy of ipTM scores has been debated and therefore, we tested other scoring methods such as AlphaBridge and ipSAE (Álvarez-Salmoral *et al*, 2026; Dunbrack Jr, 2025) (Fig. S8 and extended data 1) but also designed our own scoring model where we specifically looked at the average ipTM score between each of the five VirB6-like proteins and its nearest 2 TMH protein (see Fig. 3B, Fig. S8 and extended data 1). This scoring method, which we named chain-pair ipTM, allowed a better separation between the models and aided in identifying reliable VirB2-like candidates (Fig. 3B).

Based on visual inspection of the models and the distribution of the scores for candidate VirB2-like pili models and the other 2 TMH protein models, we choose an ipTM cut-off score of 0.7 and a chain-pair ipTM cut-off score of 0.65 as lower cut-off values for a positive model. These are conservative values that do exclude 3 of the 14 VirB2-candidates (indicated in grey in Fig. 3A-B) to avoid false negatives.

All selected candidates for VirB2-like proteins for the reference plasmids were predicted to have an outside-inside-outside orientation in the membrane, with or without a signal peptide (extended data 1), just as PrgF_B2_ and all characterized VirB2-like proteins from G-systems (Fig. S1). This orientation, which excludes VirB3-like proteins, was therefore used as an additional criterium when screening the representative dataset from 172 plasmids for VirB2-like candidates. Using these criteria (50-170 amino acids and 2 predicted TMH in the expected orientation), potential VirB2-like candidates were found in all plasmids. These candidates were all modelled together with their respective VirB6-like protein as described earlier. We find reliable, pilus-like models for ∼70% of our representative plasmids (Fig. 3E-F, 4A and Fig. S9, extended data 1), with 120/172 with an ipTM score of at least 0.7 and 124/172 with a chain-pair ipTM score of at least 0.65.

The identified VirB2-like candidates were found to be encoded in close genetic proximity to the corresponding *virB6* gene (Fig. S10, extended data 1). Only 4 plasmids from our dataset had multiple *virB2* or *virB6* genes (extended data 1).

Noteworthy examples of G+ T4SSs for which AlphaFold could confidently model a pilus-like structure include plasmids from the important clinical pathogens *Clostridium perfringens* (pCW3), *Staphylococcus aureus* (pSK41), *Clostridium botulinum* (in the neurotoxin gene cluster on pCLJ) as well as pIP501, one of the few other characterized G+ T4SSs (Thompson & Collins, 2003; Kurenbach *et al*, 2003) (Fig. 3 A-B & 4B). We measured the diameter of the pili lumen formed by the VirB2-like protein in these models and although the measurements should be regarded as approximates, they range from ca 14 Å in pCF10 (Fig. 1D) up to ca 45 Å in pCW3 (Fig. 4B). The modelled surface charge of the pili lumen in these models vary, with pCW3 being predominantly positive, pCLJ having a mixed charge distribution, while pSK41 and pIP501 having a more neutral and negative surface charge (Fig. S11).

**Figure 4.**
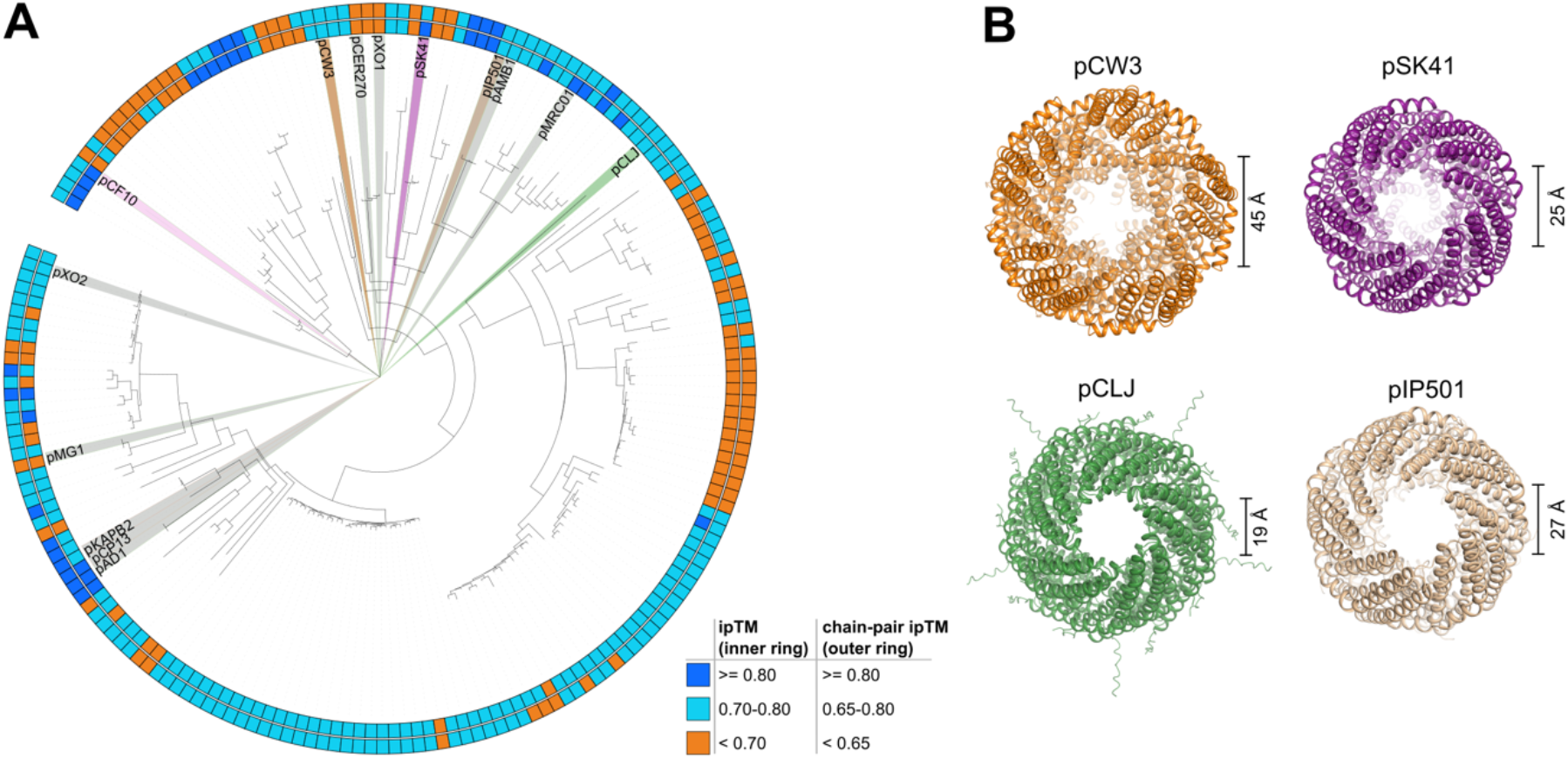
Bioinformatics analysis of VirB2-like proteins in G+ T4SSs. A) Phylogenetic tree of the VirB6-like proteins of G+ T4SSs, showing the representative classes and the 14 reference plasmids. Each coloured box indicates the score when 10 VirB2- and 5 VirB6-like proteins are modelled using AlphaFold3 by ipTM (inner ring) and chain-pair ipTM (outer ring) score. B) Top views of selected VirB2-VirB6-like protein complexes from *C. perfringens* (pCW3, ipTM: 0.75, chain-pair ipTM: 0.76), *S. aureus* (pSK41, ipTM: 0.81, chain-pair ipTM: 0.76), *C. botulinum* (pCLJ, ipTM:0.76, chain-pair ipTM: 0.70), and *E. faecalis* pIP501 (ipTM: 0.75, chain-pair ipTM: 0.78).

Overall, this bioinformatics analysis indicates that VirB2-like proteins are likely to be present in the vast majority of G+ T4SSs and that they form a pilus-like structure together with their corresponding VirB6.

## Discussion

Gram-positive T4SSs have long been described to lack conjugative pili. Competence pili for the uptake of DNA do exist in G+ bacteria(Chen *et al*, 2006; Lam *et al*, 2021), but these are not found in T4SS operons and differ greatly from conjugative VirB2-like pili. Since no previously described G+ T4SS component could form a long enough scaffold to cross the cell-wall of even just the donor cells, we wondered if VirB2-like pili forming proteins could have been overlooked. AlphaFold provided high-confidence models of PrgF_B2_, from pCF10, forming a pilus-like structure together with PrgH_B6_. Importantly, two experimental structures of VirB2-VirB6 complexes have just recently been resolved, namely from the Gram-negative pKM101 and R388 systems(Bloch *et al*, 2026; Paillard *et al*, 2026). Our modelled PrgF_B2_-PrgH_B6_ complex, as well as the other VirB2- and VirB6-like complexes, strongly resemble the experimental data from these Gram-negative systems.

We confirmed the PrgF_B2_-PrgH_B6_ pilus model by combining mutagenesis and *in vivo* assays and showed that PrgF_B2_ is essential for conjugation. As far as we are aware this is the first VirB2-like protein identified in a Gram-positive T4SS, but by using a combination of recently developed bioinformatics software, we identified VirB2-like proteins in the majority of G+ T4SSs. Although we recognize the limitations of AlphaFold modelling, our conservative cut-off scores effectively distinguished VirB2-like proteins from other 2 TMH proteins with similar length (Fig. 3A-B, Fig. S3, Table S2) and identify just a single VirB2-like candidate out of all open reading frames for ∼70% of the plasmids (Extended data 1). We propose that the newly identified VirB2-like proteins provide the missing substrate passageway through the thick cell-wall of G+ bacteria, by assembly with their VirB6-like protein as has just recently been shown for the G-systems R388 and pKM101(Paillard *et al*, 2026; Bloch *et al*, 2026).

To our knowledge, no conjugative pilus has been visualized using electron microscopy on T4SS expressing G+ bacteria. Possibly, the thick and heavily crosslinked cell wall obscures pili-like features and/or pili extension occurs only transiently, when a mating pair has formed and does not extend far outside the cell. As adhesins mediate close cell-to-cell contacts in many G+ systems, we hypothesize that the role of the pili would be to cross through the cell-wall layers of both donor and recipient cells. This would be reminiscent of G-systems with dynamic pili, such as the F-plasmid, when they are in their retracted state (Margaret Clarke *et al*, 2008). We have extensively tried to study the pCF10 T4SS and its conjugative pilus using both fluorescence and electron microscopy. However, due to very low endogenous expression levels this has proved to be extremely challenging.

We searched but could not identify any VirB5-like protein to act as a pilus tip in pCF10. This does not conclusively prove that no VirB5-like proteins exist, but it is worth noting that there also are G-systems without identified pili tips, such as the T4SSs from the F-plasmid from *E. coli,* and from Ptl from *B. pertussis*, GGI from *N. gonorrhoeae* and ComB from *H. pylori* ComB (Fernandez-Gonzalez & Backert, 2014; Hamilton *et al*, 2005; Alvarez-Martinez & Christie, 2009).

In our AlphaFold models, the diameters of the modelled pili lumen vary from 14 up to 45 Å (Fig. 3B). Although these numbers should be seen as approximations, they are in line with experimentally determined pili structures from G-conjugative plasmids, many of which have a diameter of around 25 Å (Beltran *et al*, 2023). In all cases, the pore sizes of the channels are large enough to support ssDNA transfer, which has been shown to need a diameter of at least 10 Å (Heng *et al*, 2006).

Many VirB2-like proteins in conjugative pili from G-systems have been shown to bind specific lipids, usually phosphatidyl glycerol (Costa *et al*, 2016; Vadakkepat *et al*, 2024a). These lipids line the lumen and modulate the generally positive surface charge to moderately negative to facilitate efficient DNA transfer through the pili. In our modelled G+ PrgF_B2_ pilus the lumen is already negatively charged (Fig. 2C), as is the lumen of the modelled pIP501 and pSK41 (Fig. S11). This suggests that no lipid binding is necessary for these examples as has been shown for the G-RP4 pilus (Ishimoto *et al*, 2025a). iHowever, other modelled pili have an overall positive lumen, so they might bind a lipid, but this remains to be determined.

To conclude, we have here shown that G+ conjugative T4SSs depend on VirB2-like proteins for which the data strongly indicates that they form conjugative pili. Due to almost non-existing sequence homology, these proteins were overlooked until now but using newly developed bioinformatics tools we could identify them. T4SSs and VirB2-like proteins are present in numerous plasmids from a wide range of species, including those prominent in nosocomial infections, such as *E. faecium* and *S. aureus*. These findings open completely new ways to study the spread of both virulence factors and antibiotic resistance in Gram-positive bacteria. Understanding pilus regulation and biogenesis, as well as visualizing the pili remain intriguing, but challenging, questions to address in future studies.

## Methods

### In silico analysis

AlphaFold 3 modelling for PrgH**_B6_** and PrgF_B2_ and negative control runs with PrgI_B3_ and PcfH were done on the AlphaFold server (Abramson *et al*, 2024). The topology of PrgF_B2_ was predicted using DeepTMHMM (v1.0.24) (Hallgren *et al*, 2022). Surface charge calculations of the PrgF_B2_-PrgH_B6_ and other pili models were done in PyMOL (v3.1.5.1), using the APBS plugin (v3.4.1) (Jurrus *et al*, 2018). All AlphaBridge interfaces were calculated at the default confidence level.

### Construction of OG1RF pCF10ΔprgF_B2_

The construction of *E. faecalis* OG1RF:pCF10Δ*prgF*_B2_ strain was carried out with allelic exchange and counterselection aided by a pCJK218-derived plasmid (Vesić & Kristich, 2013b). The first 5 and last 8 amino acids of PrgF_B2_ were kept intact to prevent the disruption of downstream genes. 900 bp upstream and downstream regions of *prgF***_B2_** were PCR amplified from pCF10 with primer pairs of KOprgF_UF_fwd_BamHI, KOprgF_UF_rev_XbaI and KOprgF_DF_fwd_XbaI, KOprgF_DF_rev_NcoI respectively (Table S4). The PCR products were digested with either BamHI/XbaI or XbaI/NcoI and ligated into BamHI/NcoI/XbaI digested pCJK218. For the rest of the procedure the protocol from Jäger et al. (Jäger *et al*, 2022a) was followed.

### Construction of pMSP:*prgF_B2_* (wild type and mutants)

*prgF*_B2_ was PCR amplified from pCF10 with primer pairs pMSP_prgF_fwd_NcoI and pMSP_prgF_rev_XbaI (Table S4). The PCR product was digested with NcoI/XbaI and ligated into the NcoI/XbaI digested pMSP3545S:MCS vector (further called pMSP) (Sun *et al*, 2023a). After transformation and selection in *E. coli* Top10 cells, the obtained pMSP:*prgF*_B2_ vector was sequenced and used as a template for iPCR reactions to construct pMSP:prgF_D37C, pMSP:prgF_D37N, pMSP:prgF_D38C, pMSP:prgF_D38N, pMSP:prgF_R40C, pMSP:prgF_R40E, pMSP:prgF_D37NandD38N (see Table S2 for used primers). prgF-6His, prgF-HA, prgF-StrepII, prgF_T5C, prgF_T9C, prgF_S35C, prgF_T68C were instead ordered synthetically with a NcoI site around the start codon and a BamHI site after the stop codon and digested and ligated into the pMSP vector (see Table S4 for further details).

All obtained constructs were sequenced and transformed to OG1RF:pCF10Δ*prgF*_B2_ according to the procedure described in Sun et al (Sun *et al*, 2023a). Cysteine mutations, T5C, T9C, S35C, T68C were designed to be mild mutations accessible for labelling, based on the AlphaFold model and NetSurP 3.0 (Høie *et al*, 2022) scores as described in Ellison et al. (Ellison *et al*, 2019).

### Conjugation assays

Donor (OG1RF pCF10 and derivatives thereof) and recipient (OG1ES) cells were inoculated from glycerol stock and grown overnight (∼16 hrs) at 37 °C and 180 rpm in BHI with the appropriate antibiotics; 10 μg/mL tetracycline (to select for pCF10) and 100 μg/mL erythromycin for donors with pMSP or for the recipient cells just with 20 μg/mL erythromycin (see strains and plasmid list Table S4). The following day the cultures were refreshed in BHI without antibiotics and incubated for 2.5 hrs at 37 °C and 180 rpm to get the cells in log-phase with the donor cells OD_600nm_ ∼ 0.3 and the recipient culture (single culture used for all donors in one experiment) OD_600nm_ ∼ 0.5. For conjugation assays with pMSP containing donors, 100 ng/mL nisin (Sigma) was added to the recipient culture just before setting up the conjugation mixtures. 1 mL of each donor culture was mixed with 1 mL of the recipient culture. Conjugation mixtures were then incubated for 2.5 hrs at 37 °C and 105 rpm. After this incubation the mixtures were serially diluted in PBS and 10 µL drops of undiluted to 10^5^ times diluted cultures were plated out in triplicates on BHI agar plates supplemented with 10 μg/mL tetracycline and 25 μg/mL fusidic acid (to select for donor cells) or 10 μg/mL tetracycline, 20 μg/mL erythromycin and 1 mg/mL streptomycin (to select for transconjugants). Plates were incubated at 37 °C for ∼42 hrs, counted and enumerated for colony-forming units (CFU). The plasmid transfer rate was determined as CFU of transconjugant over CFU of donor (T/D). To determine the background level of our OG1RF and OG1ES strains under the tested conditions, OG1RF without pCF10 was also used as a negative control “donor”, where the CFU of “donor” was based on BHI agar plates with only fusidic acid.

To be able to label cysteine variants of PrgF_B2_ in the presence of the rest of the T4SS before conjugation the protocol was adjusted. The overnight cultures of donors and recipient were refreshed 1:100 and incubated for 2 hrs at 37 °C and 180 rpm. Then not only nisin, but also cCF10 was added to the donor cultures at a final concentration of 50 ng/mL and 10 ng/mL respectively and the agitation was reduced to 105 rpm for all cultures. After 1 hour of induction, 500 μL aliquots for each donor culture were spun down at 16,000 rcf for 1 minute. Cell pellets were washed in 1 mL PBS, spun down again and resuspended in 250 μL PBS. Then two 100 uL aliquots were taken out to which either 1 μL 125 mM maleimide-PEG11-biotin (EZ-link) in DMSO was added (for PEG-labelling) or 1 μL DMSO (for control). All aliquots were mixed by vortexing and incubated for 10 minutes at 25 °C. Afterwards 200 μL of the recipient cell culture (in BHI, without nisin or cCF10) was added to each of the (labelled or control) donor fractions and mixed by vortexing. These conjugation mixtures were incubated statically at 37 °C for just 15 minutes. The rest of the procedure was done as described above.

All conjugation data shown in figure 2 was done with strains with a pMSP plasmid, either empty (first two bars of Fig. 2A) or with *prgF*_B2_ (all other bars as indicated) and with nisin induction as specified above. The conjugation data shown in figure S5 is from strains without pMSP vector and without the addition of nisin. All conjugation results are shown as bars of the mean, with error bars showing the standard error of the mean of at least three biological replicates, that are also shown as individual datapoints (black dots). To include data with no detected transconjugants in the analysis, these were given a very small score, 10^-6^, therefore indicated as ND = not detected on the Y-axes. Grey area indicates the background T/D value as calculated from conjugation experiments of OG1RF without pCF10 (Fig. S5). Data visualization and analysis was done with GraphPad Prism (version 11, GraphPad software).

### Cell culture, protein expression and membrane preparation

*E. faecalis* OG1RF strains containing either wild-type pCF10 and pMSP (empty vector), or pCF10Δ*prgF*_B2_ and pMSP:*prgF*_B2_-6xHis, were inoculated from glycerol stock and grown overnight at 37 °C and 180 rpm in tryptic soy broth (TSB; Merck) with 10 µg/mL tetracycline and 100 μg/mL erythromycin. The next day, overnights were used in a 1:40 ratio to inoculate 2 L of TSB with 10 µg/mL tetracycline and grown until an OD_600nm_ of approximately 0.8 (37 °C, 180 rpm). Shaking was then reduced to 80 rpm, and expression of both plasmids was done with 10 ng/mL cCF10 and/or a 1:1000 dilution of nisin-containing supernatant from *Lactococcus lactis* NZ9100 cultures as indicated. After 120 minutes of induction, cells were harvested by centrifugation (4,000 x *g*, 10 min), washed once with membrane buffer (50 mM potassium phosphate buffer (KPi), 150 mM NaCl, with 10% glycerol, pH 7.4) and flash-frozen in lN_2_ before storage at -20 °C.

For membrane preparation, cell pellets from 2 L of cultures were resuspended in 80 mL ice-cold membrane buffer, supplemented with 1 mM MgSO_4,_ 0.2 mM 4-(2-aminoethyl)benzenesulfonyl fluoride hydrochloride (AEBSF; ITW Reagents) and a few mg of DNAse I (BioRad). Resuspensions were lysed using a Cell Disruptor (Constant Systems) for 3 passes at 40 k psi. Lysates were pelleted to remove debris (13,000 x *g*, 30 minutes, 4 °C) and the supernatant was further processed by ultracentrifugation (60,000 x *g*, 1 hour 15 minutes, 4 °C) to retrieve the membrane fraction. Membranes were resuspended in 1.5 mL of membrane buffer, divided into 0.5 mL aliquots, flash-frozen in lN_2_ and stored at -80 °C until use.

#### Cobalt IMAC pulldown of PrgF_B2_-6xHis

For each purification a single 0.5 mL membrane aliquot was rapidly thawed and solubilised on ice in 20 mM KPi (pH 7.0), 150 mM NaCl, 10% glycerol, 1% lauryl maltose neopentyl glycol (LMNG; Anatrace) and 1 mM AEBSF with a total volume of 2 mL. After 1 h of gentle rotation at 4°C, the solubilised membranes were ultra-centrifugated to remove debris (30,000 x *g*, 30 minutes, 4 °C) and the supernatant was incubated with 250 µL pre-equilibrated Cobalt-nitriloacetic acid agarose beads (Invitrogen) for 1 hour at 4 °C in the presence of 10 mM imidazole (added from 1M stock at pH 7.5). Afterwards, the mixture was loaded into a small gravity flow column (BioRad) and washed with 60 column volumes (CV) 50 mM HEPES (pH 7.5), 150 mM NaCl, 0.003% LMNG, 10 mM imidazole (pH 7.5) and 0.2 mM AEBSF. Elution was done in two steps with 250 µL 50 mM HEPES (pH 7.5), 150 mM NaCl, 0.003% LMNG, 150 mM imidazole (pH 7.5) and 0.2 mM AEBSF with 5 minutes incubation (at 4°C) each time. The two elution fractions were then pooled and concentrated to 60 µL using a 3 kDa cutoff filter (Amicon Ultra). Glycerol was added to 20% and samples were flash-frozen in lN_2_ and stored at -80 °C until use for SDS-PAGE.

#### SDS-PAGE and Western blotting

The total protein concentration in the membranes was assayed by the bicinchoninic acid method (Pierce kit, Invitrogen). Membranes were diluted to the indicated concentration and SDS sample buffer with β-mercaptoethanol was added. To check the expression levels of the downstream genes a 30 µL membrane fraction with 20 µg of total protein, for both pCF10 and pCF10Δ*prgF_B2_*, was loaded onto Novex Tris-Glycine Mini Protein Gels, 4–12%, 1.0 mm, WedgeWell (Invitrogen) and run at 225 V for 40 minutes, or until the dye front nearly reached the bottom of the gel. To detect PrgF_B2_-His, 5 µL membrane fractions with 100 µg of total protein and 20 uL of the concentrated elution fractions, all in SDS sample buffer with β-mercaptoethanol, were loaded onto 4% stacking and 16% resolving Shägger gels (Schägger, 2006).

Transfer to 0.2 µm (PrgF_B2_-His) or 0.44 µm (downstream genes expression) PVDF membrane (Amersham) was done by semi-dry transfer at 25 V, 1.3 A for 5 min (PrgF_B2_-His) or 7 min (downstream genes expression) with the TransBlot Turbo (BioRad). Blots were blocked in 2% skim milk powder in PBS with 0.1% Tween-20 (PBS-T) for 1 hr, washed 3 times in PBS-T and probed in PBS-T with 2% milk and one of the following primary antibodies: anti-His (1:1000, Invitrogen), anti-PrgL (1:1000 custom order, AgriSera) (Jäger *et al*, 2022b), anti-PcfC (1:1000) (Chen *et al*, 2008). After 1 hr incubation, the blots were again washed 3 times in PBS-T and then probed with HRP conjugated secondary antibodies: Goat anti-Mouse IgG (H&L, Agrisera) for the anti-His and Goat anti-Rabbit IgG (H&L, Agrisera) for the downstream genes. After 3 more PBS-T washes, blots were incubated for 2-5 min with ECL SuperBright substrate (Agrisera) and imaged by chemiluminescence on a ChemiDoc HP (BioRad).

### Bioinformatics analysis of G+ plasmids

Plasmid sequences from 72,360 plasmids were retrieved from PLSDB (v2024_05_31_v2) (Molano *et al*, 2025). Plasmids annotated as conjugative and from G+ origin, as identified by PLSDB, were selected creating a set of 1,787 plasmids for further analysis. All proteins from these plasmids were clustered at 90% identity and 90% coverage using MMseqs2 (v18-8cc5c) (Steinegger & Söding, 2017) resulting in 28,172 protein clusters. Protein structures for these protein representatives were predicted using ESMfold (v1.0.3, ESM-2-3B), and transmembrane helices and signal peptides were predicted using DeepTMHMM (v1.0.24) (Lin *et al*, 2023; Hallgren *et al*, 2022).

Proteins with predicted structures similar to VirB6/PrgH_B6_ were identified using Foldseek (v10-941cd33) (van Kempen *et al*, 2024), resulting in 1,113 proteins originating from 174 protein clusters. For each cluster, a representative VirB6-like protein was selected. In two cases, the representatives of two different clusters belonged to the same plasmid (as two plasmids have two instead of a single VirB6-like protein, see extended data table 1). Therefore, this clustering resulted in 172 representative plasmids that were further analysed. 14 additional references plasmids were manually selected to represent clinically important plasmids that have been studied (Abramson *et al*, 2024). For tree construction, VirB6-like proteins were aligned using Muscle5 (v5.3), sparse columns were removed using trimAl (v1.5.0) and unrooted trees were constructed using IQ-TREE3 (v3.0.1) (Edgar, 2022; Capella-Gutiérrez *et al*, 2009; Wong *et al*, 2025). Trees were then rooted at midpoint and visualised using iTOL (v7) (Letunic & Bork, 2024).

As some VirB6 proteins are too large to model with AlphaFold due to large cytosolic C-terminal domains, we decided to trim all VirB6 sequences 5 amino acids after the end of the last predicted transmembrane helix (TMH) for further analyses. To identify potential VirB2-like proteins we searched for proteins with a length between 50-170 amino acids and with a predicted DeepTMHMM topology of (S)OMIMO: with/without Signal peptide, Outside, Membrane, Inside, Membrane, Outside in each of the 186 selected plasmids. For the 14 reference plasmids all 50-170 amino acids long proteins with two TMHs were taken along, thus including proteins with an (S)IMOMI orientation, such as VirB3-like proteins. Proteins with a signal peptide were further analysed without this part of the sequence. For all 186 plasmids, each of the selected 2 TMH proteins were modelled on the AlphaFold server (Abramson *et al*, 2024) as 10 copies together with 5 copies of their corresponding VirB6-like protein.

To specifically assess the interaction score between the 2 TMH protein to evaluate and their VirB6-like protein we developed a “chain-pair ipTM” score. Each chain’s centroid was computed as the mean position of all its Cβ atoms. The five VirB2 chains whose centroids lay closest to any VirB6 centroid were selected, and each was assigned its highest chain-pair ipTM against a VirB6 chain. The used metric is the mean of those five values. AlphaBridge (GitHub commit e6ff51c) (Álvarez-Salmoral *et al*, 2026) was run locally on the highest ranked model of each AlphaFold 3 prediction. The scores were recomputed to be consistent with the rules of the AlphaBridge web server.

## Data availability

All sequences and structure models used for the bioinformatics have been deposited to Zenodo and can be found with the following link: https://doi.org/10.5281/zenodo.24173912

## Author contributions

Josy ter Beek: Conceptualization, Methodology, Investigation, Writing – Original Draft, Writing – Review & Editing. Dennis Svedberg: Methodology, Investigation, Data curation, Writing – Review & Editing. Kieran Deane-Alder: Investigation, Writing – Review & Editing. May Ibrahim: Investigation, Writing – Review & Editing. André Mateus: Validation, Writing – Review & Editing, Supervision, Funding acquisition. Ronnie P-A Berntsson: Conceptualization, Writing – Original Draft, Writing – Review & Editing, Supervision, Funding acquisition.

## Acknowledgments

The authors thank Prof. Peter J Christie for providing antibodies against PcfC. This work was supported by grants from the Swedish Research Council (2023-02423), Knut and Alice Wallenberg Foundation, Insamlingsstiftelsen för medicinsk forskning and Kempestiftelserna (SMK-2059) to R.P-A.B., from European Union’s Horizon 2020 research and innovation programme under the Marie Skłodowska-Curie grant agreement No 101202752 to K.D-A., and from Kempestiftelserna (JCK-3126) to A.M. The bioinformatics predictions made in this study were enabled by the Berzelius resource provided by the Knut and Alice Wallenberg Foundation at the National Supercomputer Centre.

## Supplementary Tables and Figures

**Table S1:**
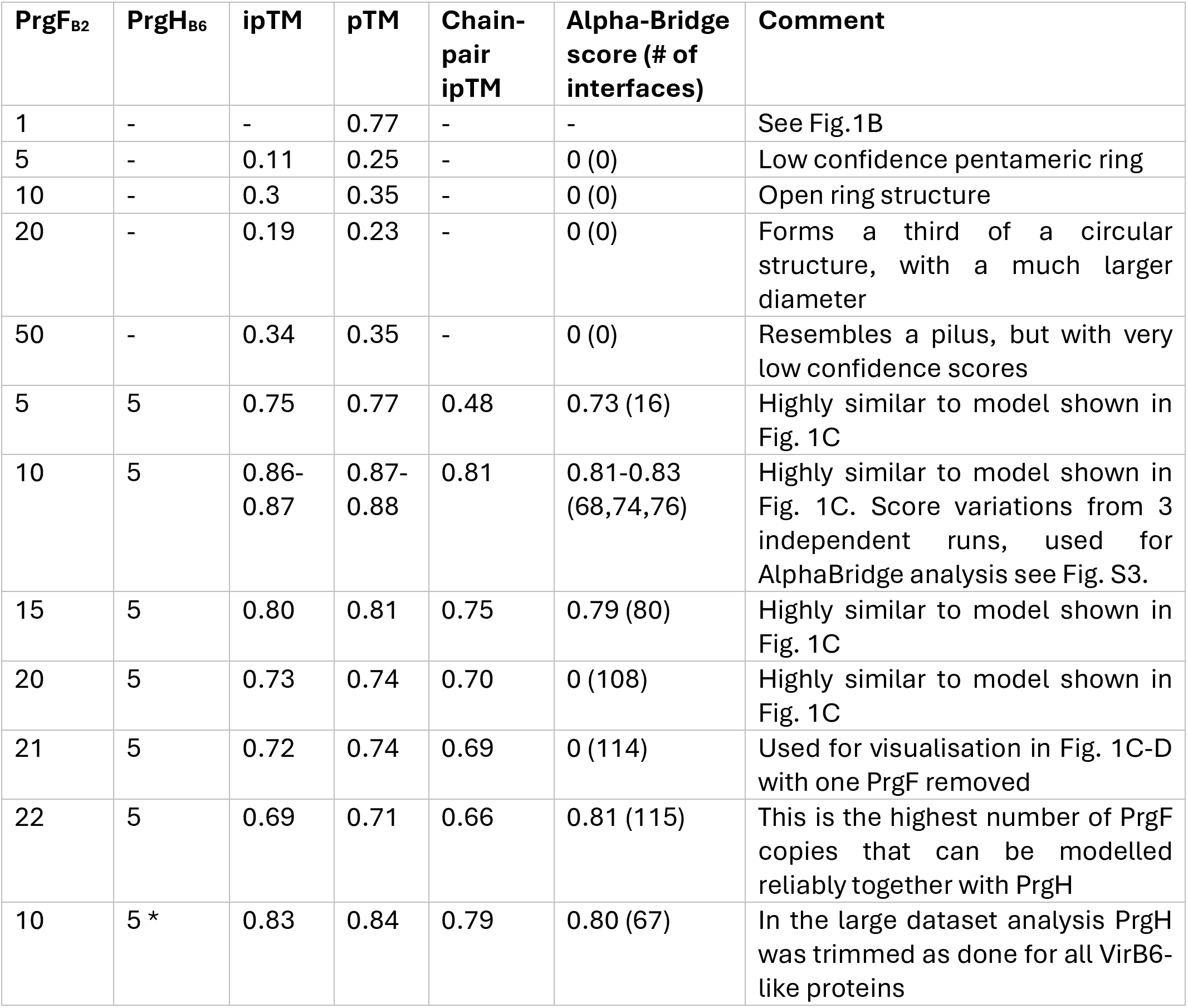
Scores and short descriptions of various AlphaFold 3 models of PrgF_B2_ with or without PrgH_B6_.

**Table S2:**
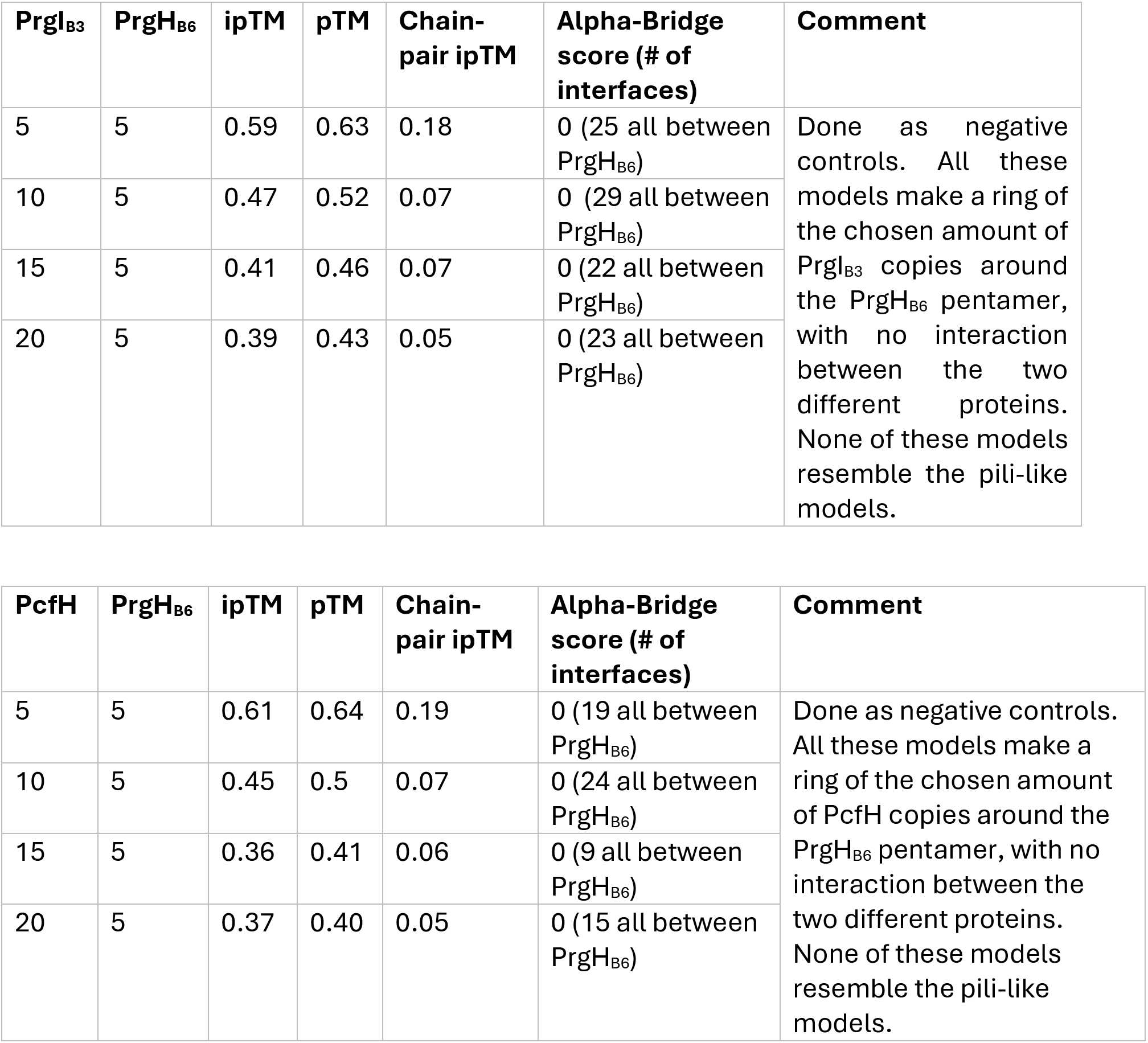
Scores and short descriptions of AlphaFold 3 models of PrgH_B6_ and PrgI_B3_ and PrgH_B6_ and PcfH, which were used as negative controls.

**Table S3.**
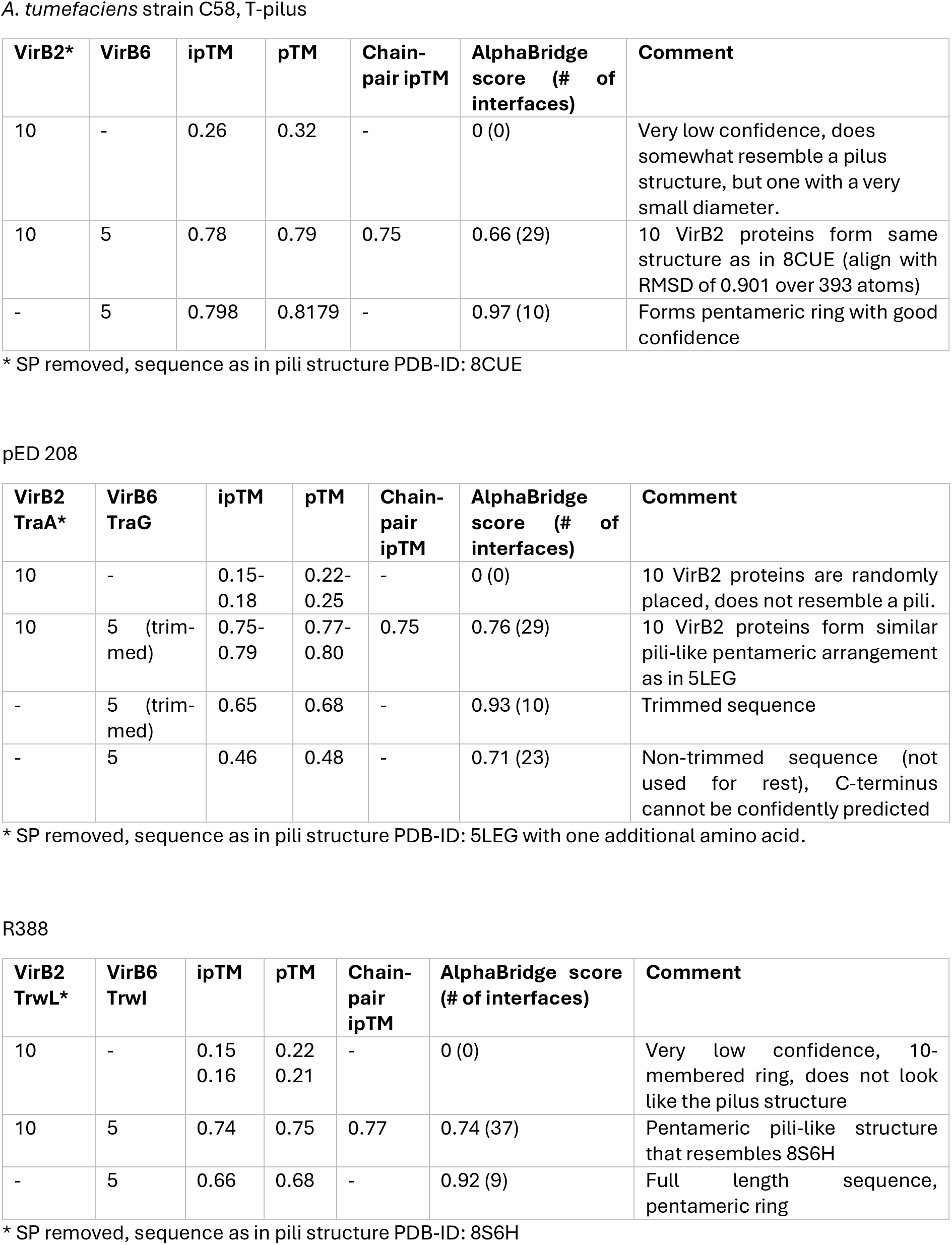
Scores and short descriptions of various AlphaFold 3 models of known pili-forming VirB2-proteins from T4SSs of Gram-negative bacteria.

| *A. tumefaciens* strain C58, T-pilus |  |  |  |  |  |  |
| **VirB2\*** | **VirB6** | **ipTM** | **pTM** | **Chain-pair ipTM** | **AlphaBridge score (# of interfaces)** | **Comment** |
|---|---|---|---|---|---|---|
| 10 | - | 0.26 | 0.32 | - | 0 (0) | Very low confidence, does somewhat resemble a pilus structure, but one with a very small diameter. |
| 10 | 5 | 0.78 | 0.79 | 0.75 | 0.66 (29) | 10 VirB2 proteins form same structure as in 8CUE (align with RMSD of 0.901 over 393 atoms) |
| - | 5 | 0.798 | 0.8179 | - | 0.97 (10) | Forms pentameric ring with good confidence |
| pED 208 |  |  |  |  |  |  |
| **VirB2<br>TraA\*** | **VirB6<br>TraG** | **ipTM** | **pTM** | **Chain-pair ipTM** | **AlphaBridge score (# of interfaces)** | **Comment** |
| 10 | - | 0.15-0.18 | 0.22-0.25 | - | 0 (0) | 10 VirB2 proteins are randomly placed, does not resemble a pili. |
| 10 | 5 (trim-med) | 0.75-0.79 | 0.77-0.80 | 0.75 | 0.76 (29) | 10 VirB2 proteins form similar pili-like pentameric arrangement as in 5LEG |
| - | 5 (trim-med) | 0.65 | 0.68 | - | 0.93 (10) | Trimmed sequence |
| - | 5 | 0.46 | 0.48 | - | 0.71 (23) | Non-trimmed sequence (not used for rest), C-terminus cannot be confidently predicted |
| R388 |  |  |  |  |  |  |
| **VirB2<br>TrwL\*** | **VirB6<br>TrwI** | **ipTM** | **pTM** | **Chain-pair ipTM** | **AlphaBridge score (# of interfaces)** | **Comment** |
| 10 | - | 0.15<br>0.16 | 0.22<br>0.21 | - | 0 (0) | Very low confidence, 10-membered ring, does not look like the pilus structure |
| 10 | 5 | 0.74 | 0.75 | 0.77 | 0.74 (37) | Pentameric pili-like structure that resembles 8S6H |
| - | 5 | 0.66 | 0.68 | - | 0.92 (9) | Full length sequence, pentameric ring |

\* SP removed, sequence as in pili structure PDB-ID: 8CUE
\* SP removed, sequence as in pili structure PDB-ID: 5LEG with one additional amino acid.
\* SP removed, sequence as in pili structure PDB-ID: 8S6H

**Table S4:**
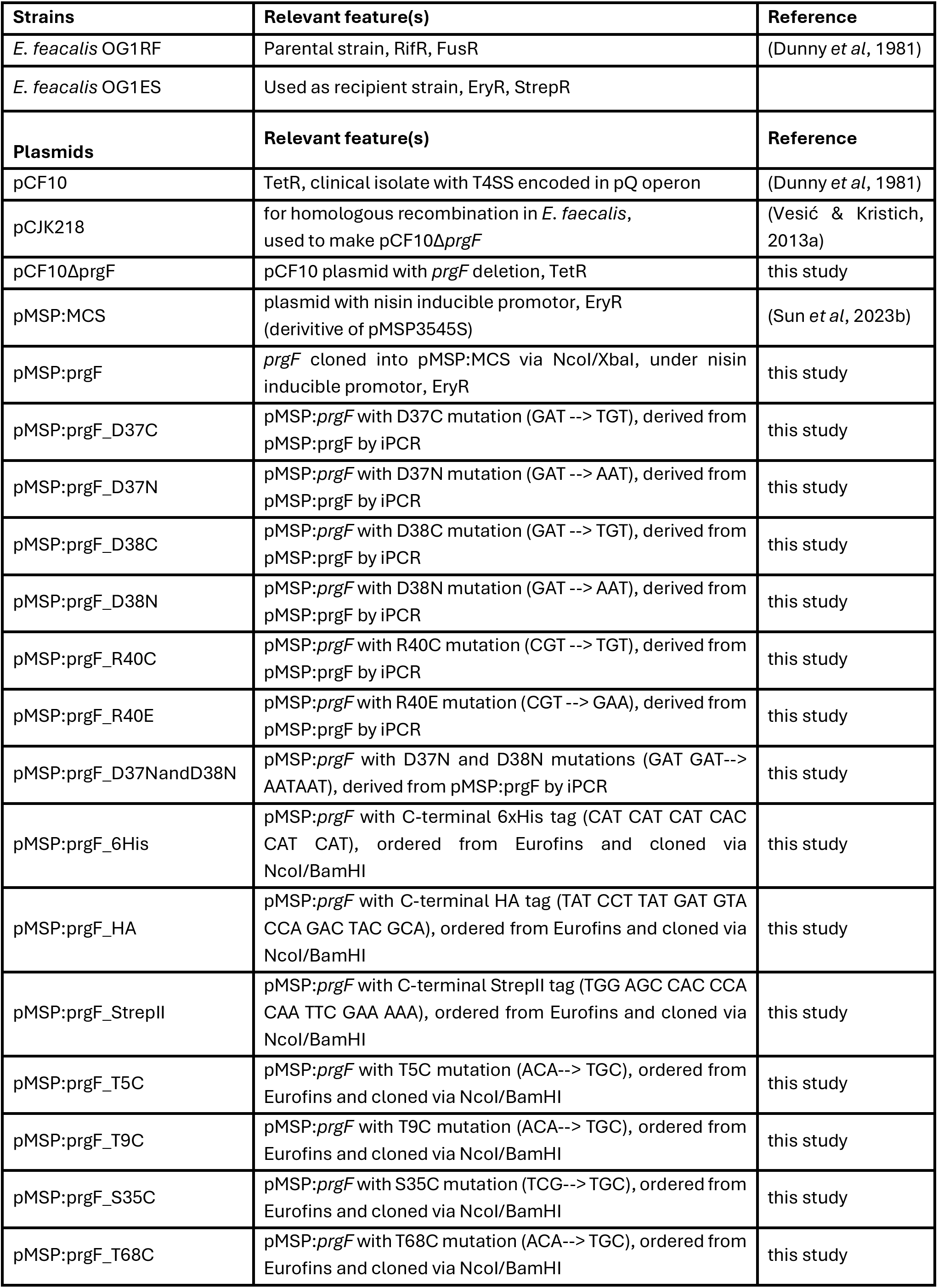
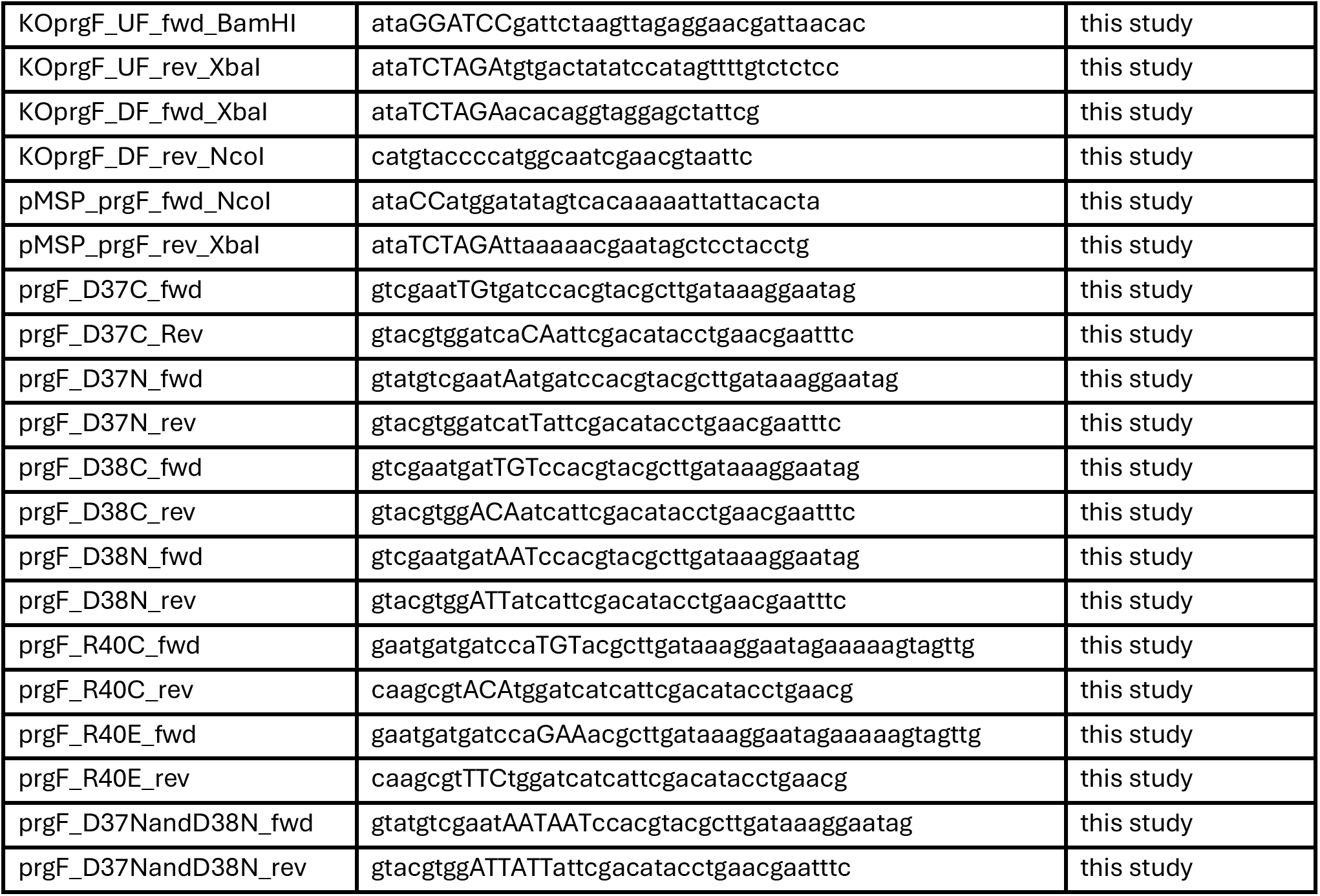
Strains, plasmids and primers used in this study.

**Fig. S1:**
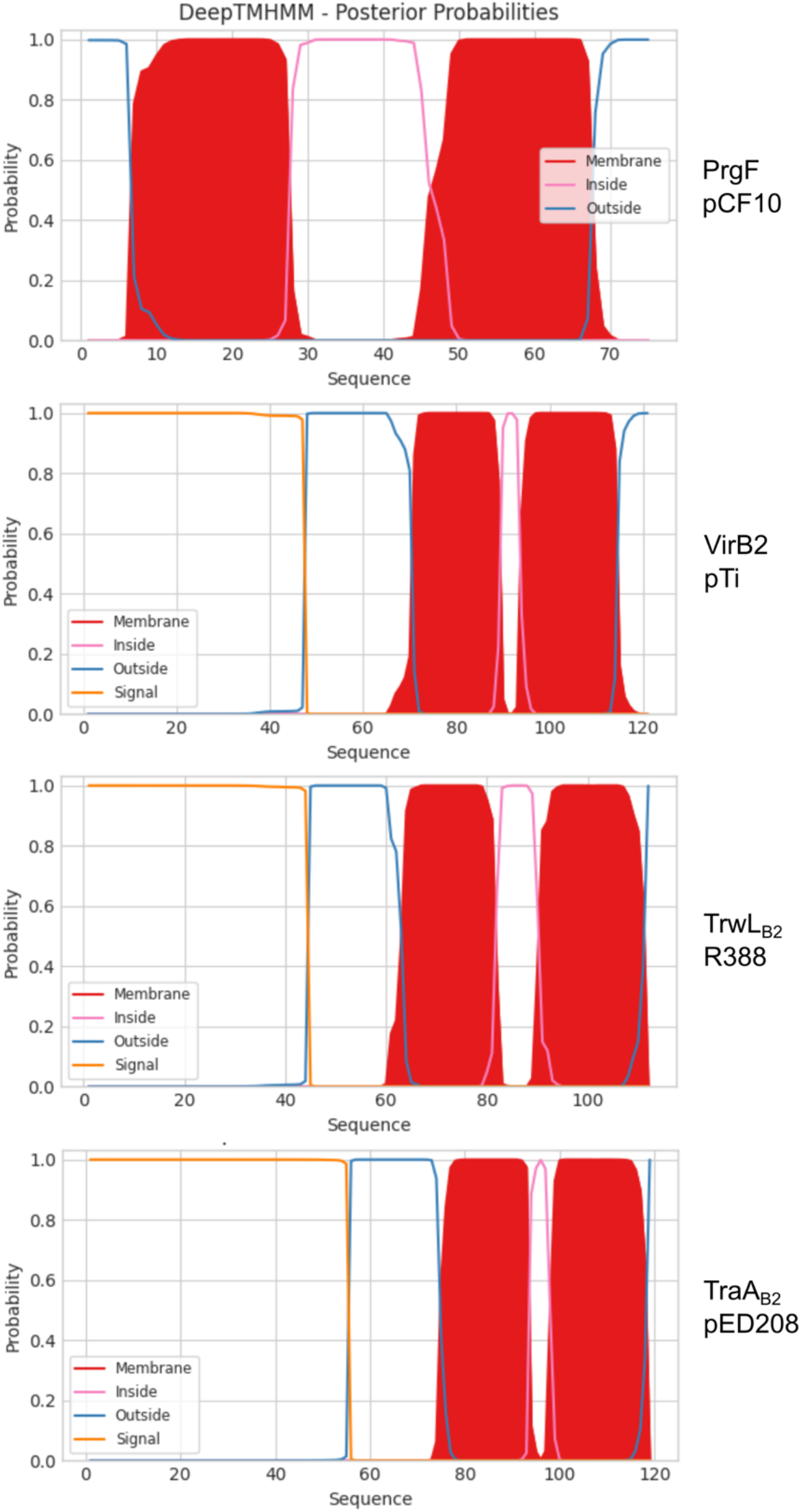
DeepTMHMM predictions of VirB2-like proteins: PrgF (pCF10), VirB2 (pTi), TrwL_B2_ (R388), TraA_B2_ (pED208).

**Figure S2.**
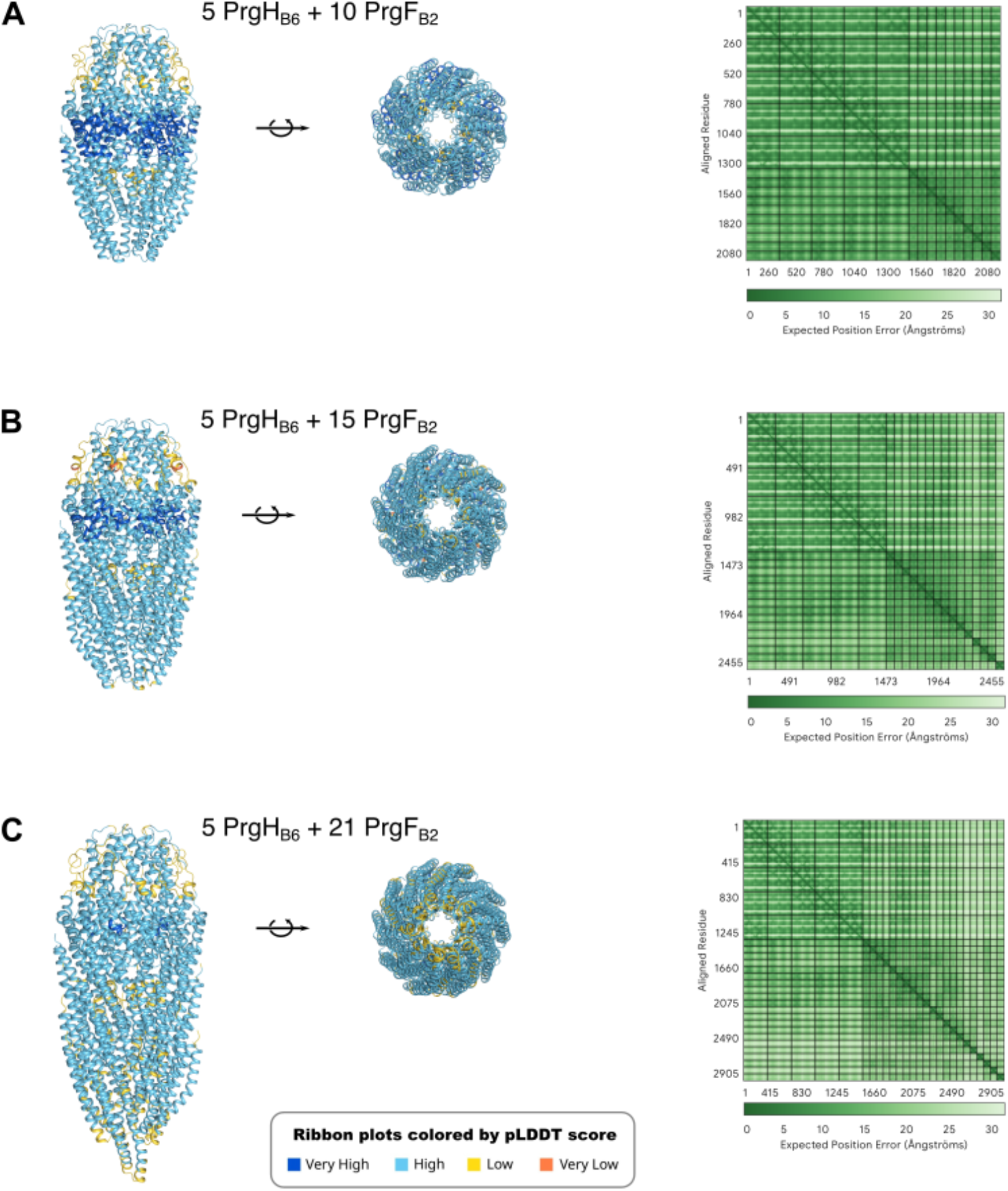
AlphaFold models of PrgF_B2_ with PrgH_B6_. All panels have 5 copies of PrgH_B6_, with increasing numbers of PrgF_B2_ (10 in panel A, 15 in panel B and 21 in panel C). Models are oriented as in Fig. 1C (side-views with PrgH_B6_ on top, and bottom-views from the PrgF_B2_-side) coloured according to their pLDDT scores, and the PAE plot of each model is shown on the right (starting with 5 copies of PrgH_B6_ from the upper left corner).

**Figure S3:**
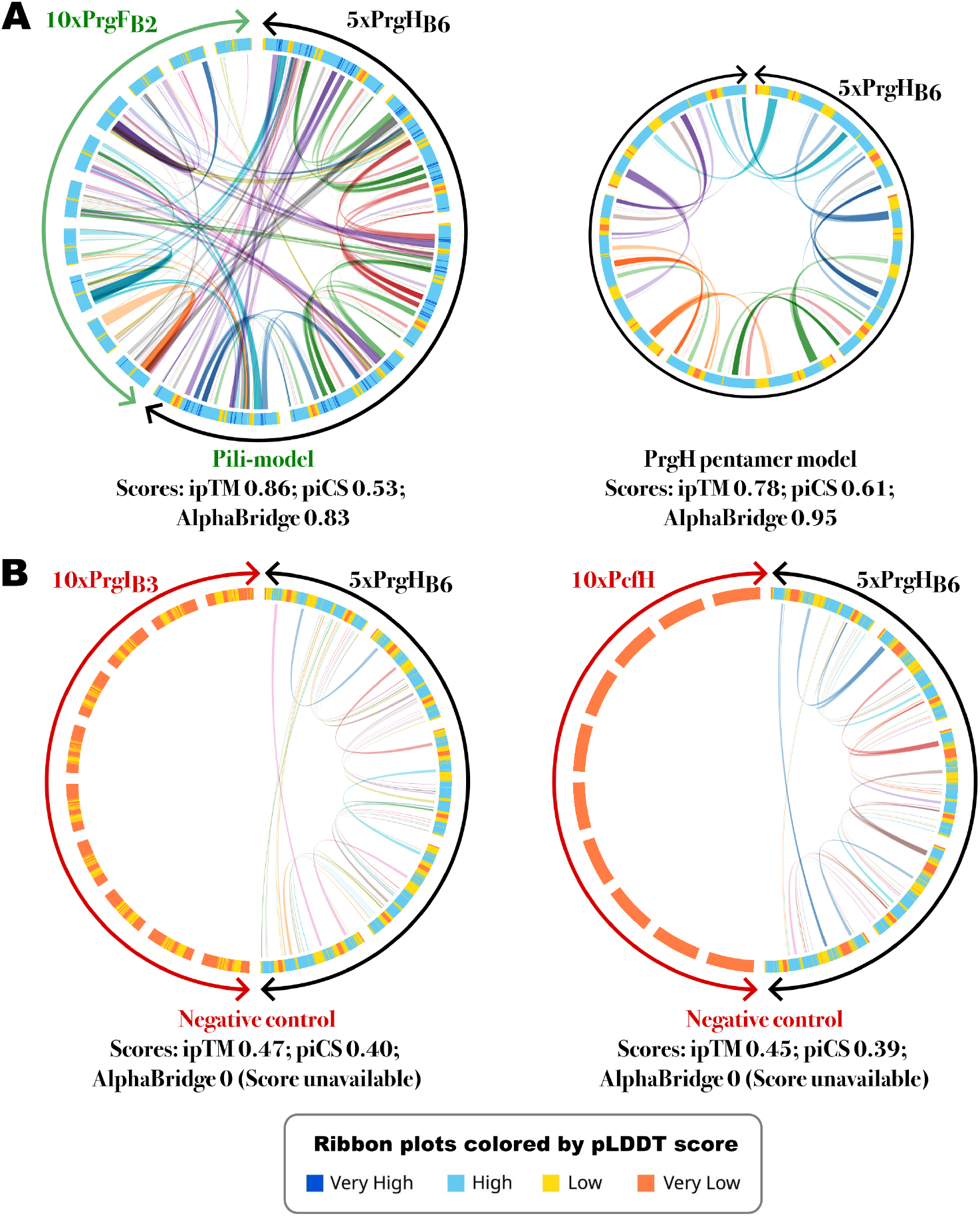
Verification of various AlphaFold models with the PrgH_B6_ pentamer by AlphaBridge at the default cutoff. A) Around 70 interactions are detected between the PrgH_B6_ and PrgF_B2_ subunits of the pili-model. B) Analysis of the PrgH_B6_ pentamer model without any additional proteins. Note that the ipTM and pLDDT scores in this model are lower than the in the pili-model shown in A, although the AlphaBridge score is higher. C and D) The negative control models with PrgI_B3_ and PcfH have much lower confidence scores and shows no interactions between these two TMH proteins and PrgH_B6_.

**Figure S4.**
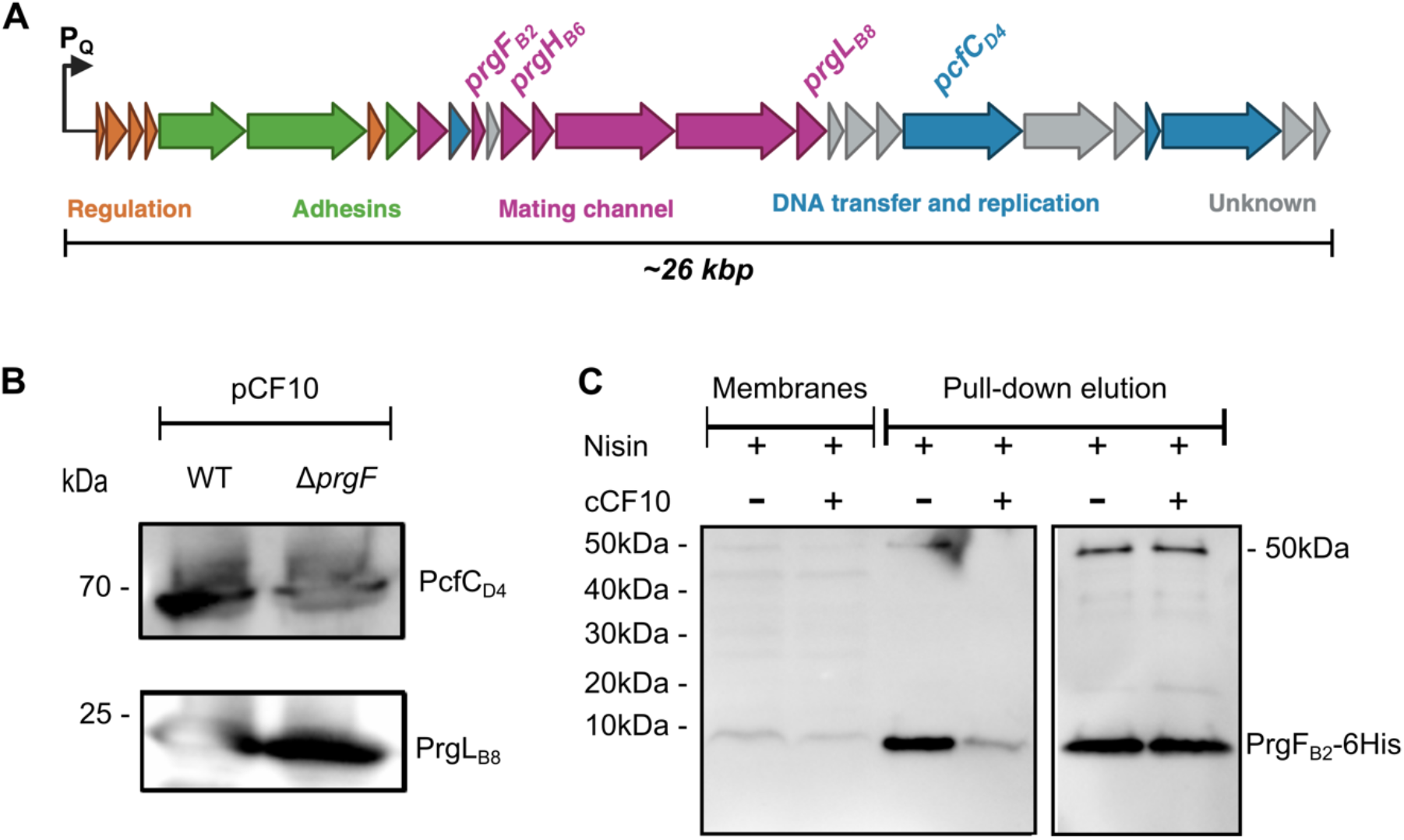
Deletion of *prgF*_B2_ does not affect the expression of downstream genes in the *prg*Q operon, PrgF_B2_ is present in cell membranes and can be pulled-down after solubilization. A) Schematic of the *prg*Q operon. B) Western blots of membrane fractions from either OG1RF with wild-type pCF10 and empty vector pMSP or OG1RF with pCF10:Δp*rgF_B2_* and pMSP:*prgF_B2_*-6xHis developed with anti-PcfC (upper pannel) or anti-PrgL (lower pannel). C) Western blot of membrane fractions and a solubilized and purified fraction from *E. faecalis* OG1RF pCF10:Δp*rgF_B2_* + pMSP:*prgF_B2_*-6xHis developed with anti-His. Cells were induced with nisin to induce PrgF*_B2_*-His expression from the nisin promotor on pMSP with or without the addition of the cCF10 pheromone which induces expression from the P_Q_ promotor.

**Figure S5:**
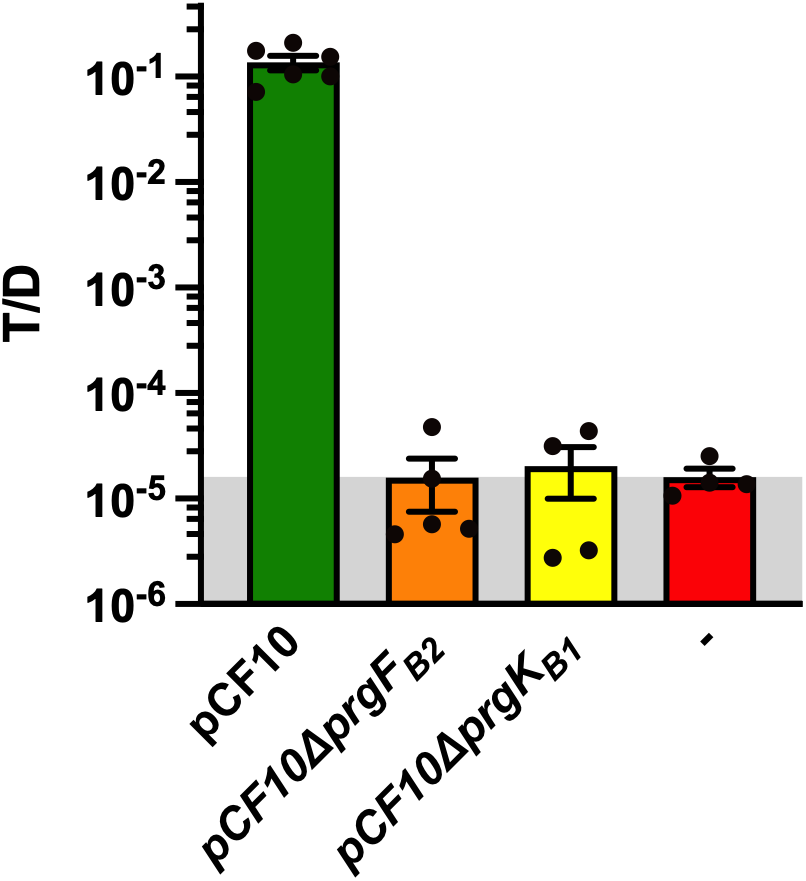
PrgF_B2_ is essential for conjugation. The OG1RF pCF10Δ*prgFB2* strain has the same conjugation background level, calculated as the number of transconjugants over donors (T/D), as the OG1RF pCF10Δ*prgKB1* strain (PrgK*B1* is known to be essential (Laverde Gomez et al, 2014)) and the same OG1RF strain without pCF10 (-). For OG1RF without pCF10 (-) the number of ”donors” were counted on a plate with just fusidic acid. All strains in this figure are without pMSP vector.

**Figure S6:**
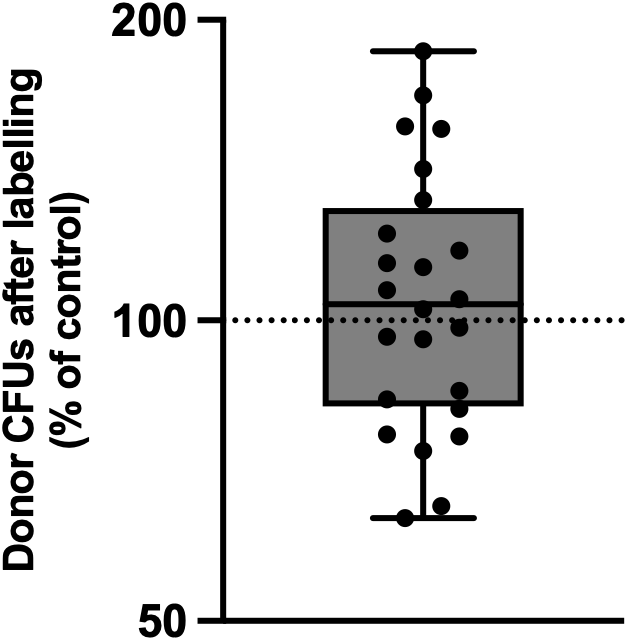
Donor cell survival is not affected by treatment with the PEG-maleimide label. Colony forming units (CFU) of PEG-labelled donor cells as a percentage of their non-labelled controls. Colony size is similar.

**Figure S7:**
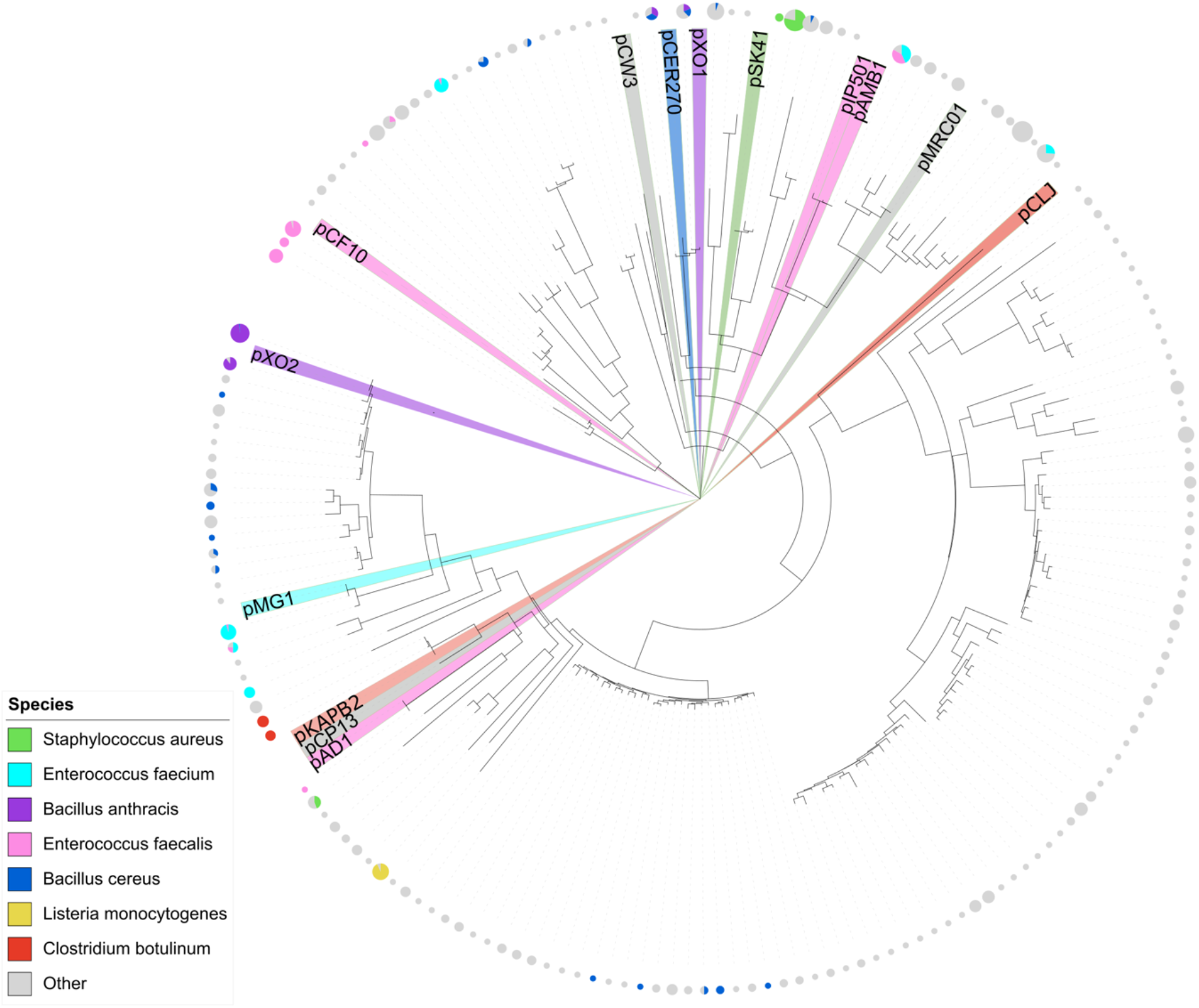
VirB6-like proteins from G+ conjugative plasmids grouped into 174 protein clusters from 172 plasmids and a wide variety of G+ species. The position of the VirB6-like proteins encoded on the 14 reference plasmids, including pCF10, are also indicated in the tree.

**Figure S8:**
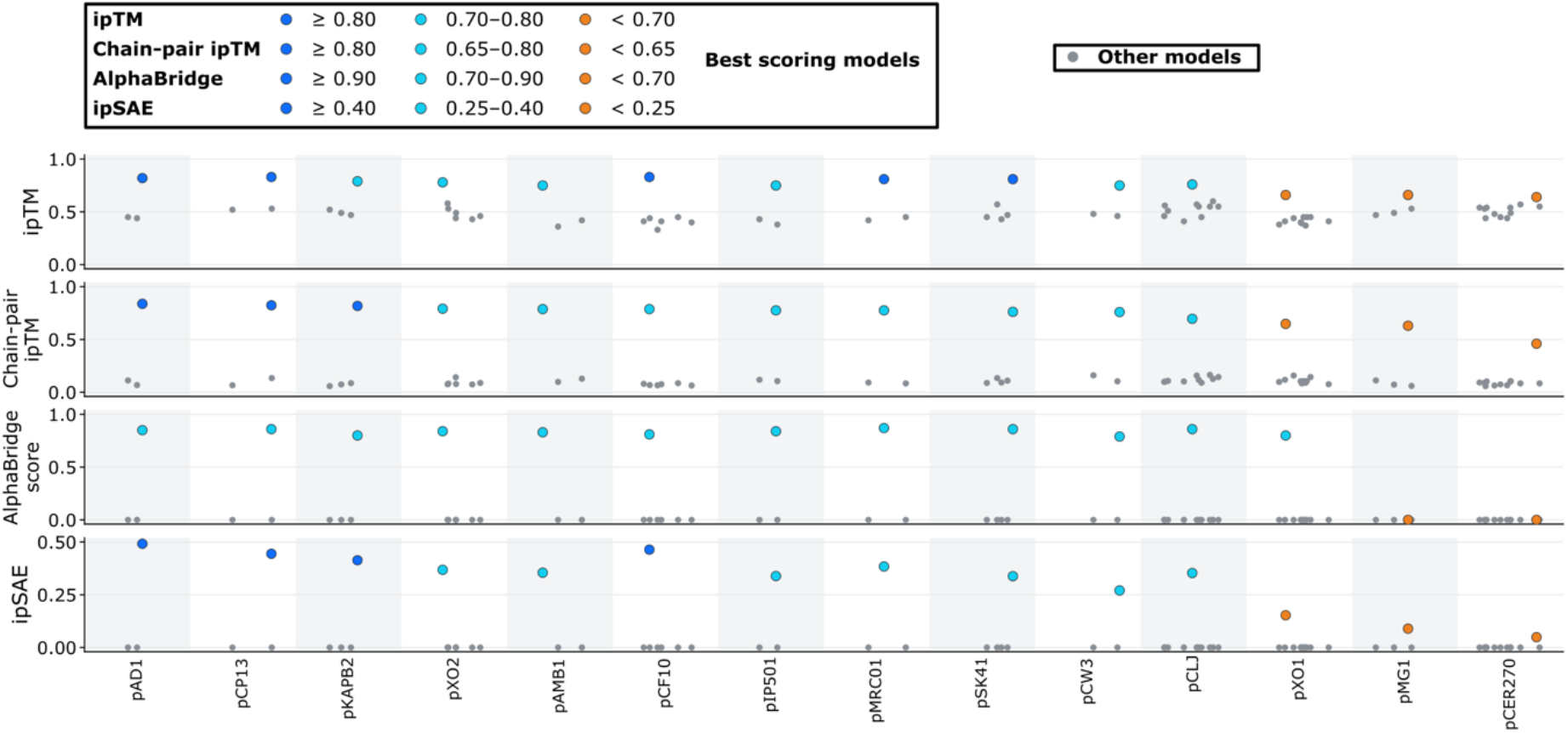
Scores of all AlphaFold models for the reference plasmids, showing from top to bottom: i) ipTM scores directly from AF3, ii) chain-pair ipTM, iii) AlphaBridge scores and iv) ipSAE score. All 2 TMH proteins modelled as 10 copies together with 5 copies of the VirB6-like proteins. VirB2-like protein candidates are highlighted in colour and were identified based on model scores and visual inspection of all models.

**Figure S9:**
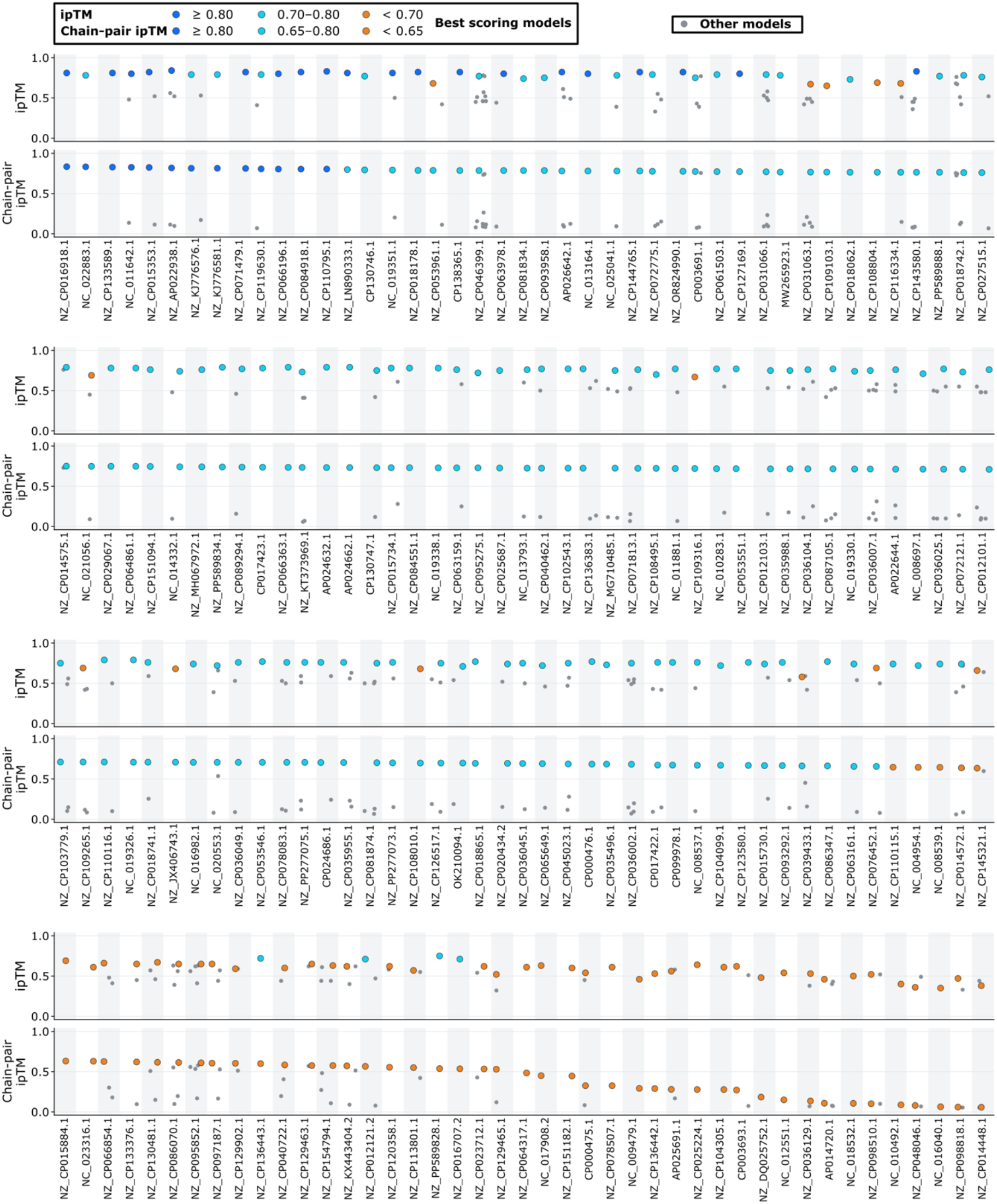
Scores as in Fig. S8 i and ii for the other 172 plasmids.

**Figure S10:**
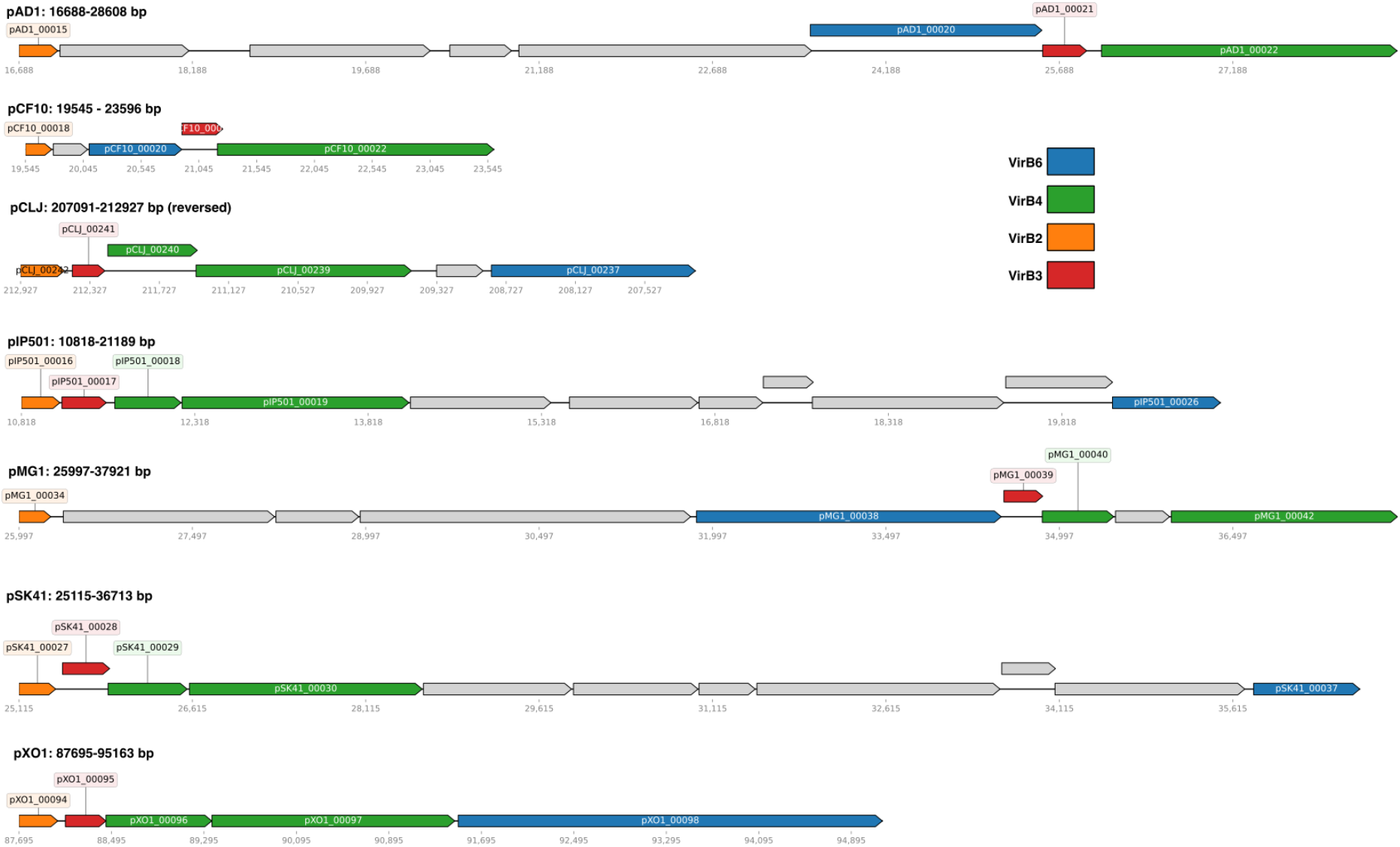
Genetic organisation of *virB2* and *virB6* genes in some of the G+ conjugative plasmids. Note that this overview only shows selected parts of the T4SS operons to visualise the location of *virB2* and *virB6*, the actual operon sizes for all these are much larger. For a complete overview of the T4SS operon of pCF10 see figure S4A.

**Figure S11.**
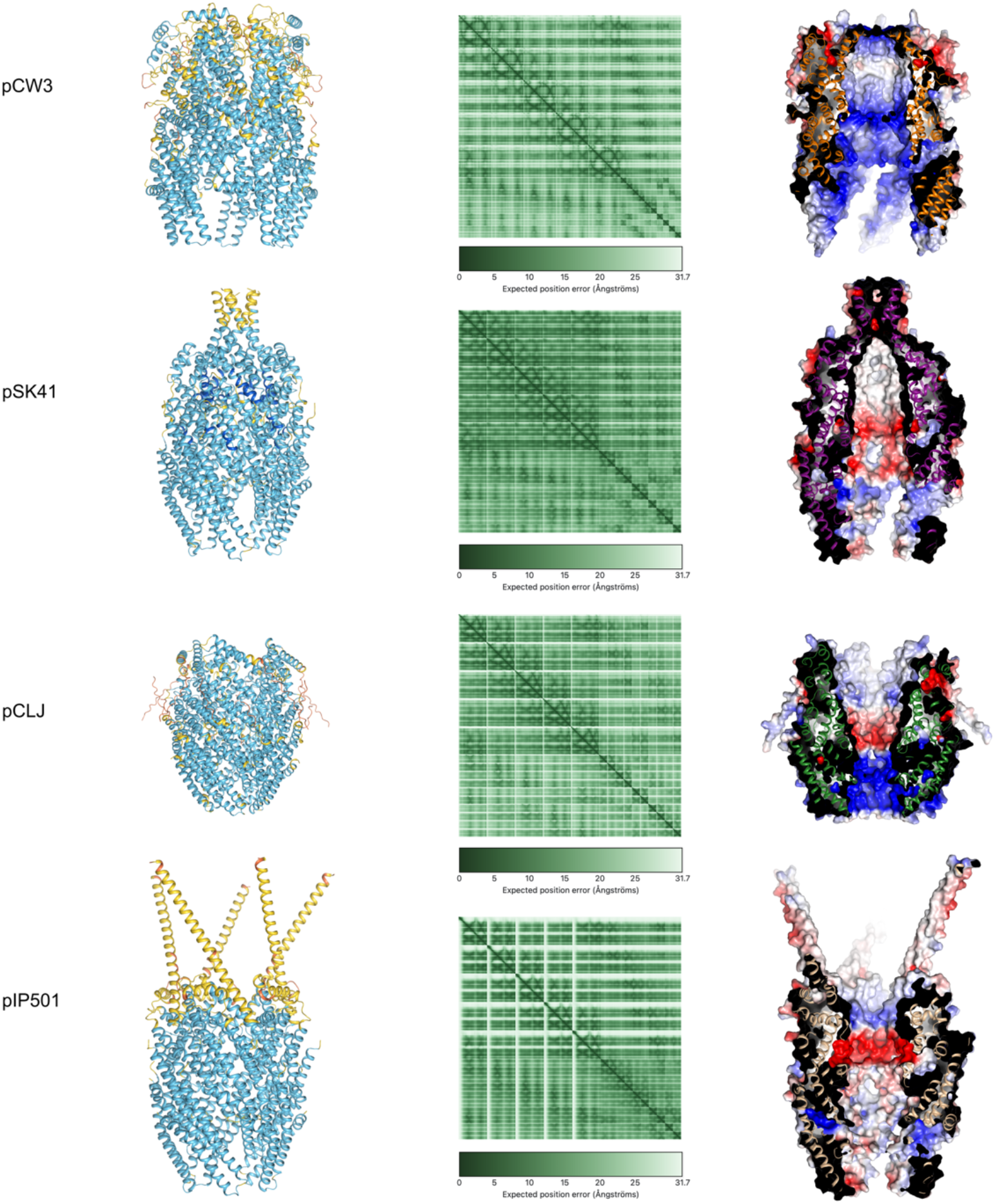
AlphaFold models of VirB2-VirB6 complexes of the T4SSs from Fig. 4B. All models have 5 VirB6-(shown at the top of each model) and 10 VirB2-like proteins (shown at the lower half of each model). In the left panel models are coloured according to their pLDDT scores, and the corresponding PAE plot of each model is shown on the middle panel. The right panels show the surface charge of the models, coloured from red (-5 kT/e) to blue (+5 kT/e).

